# Defining the seafloor microbiome of the Gulf of Mexico and its response to oil perturbation

**DOI:** 10.1101/236950

**Authors:** Will A. Overholt, Patrick Schwing, Kala M. Raz, David Hastings, David J. Hollander, Joel E. Kostka

**Author notes:** Corresponding author: Joel E. Kostka Georgia Institute of Technology 310 Ferst Drive Atlanta, Georgia 30332.

## Abstract

The microbial ecology of oligotrophic deep ocean sediments is understudied relative to their shallow counterparts, and this lack of understanding hampers our ability to predict responses to current and future perturbations. The Gulf of Mexico has experienced two of the largest accidental marine oil spills, the 1979 Ixtoc-1 blowout and the 2010 Deepwater Horizon (DWH) discharge. Here, microbial communities were characterized for 29 sites across multiple years in >700 samples. The composition of the seafloor microbiome was broadly consistent across the region and was well approximated by the overlying water depth and depth within the sediment column, while geographic distance played a limited role. Biogeographical distributions were employed to generate predictive models for over 4000 OTU that leverage easy-to-obtain geospatial variables which are linked to measured sedimentary oxygen profiles. Depth stratification and putative niche diversification are evidenced by the distribution of taxa that mediate the microbial nitrogen cycle. Further, these results demonstrate that sediments impacted by the DWH spill had returned to near baseline conditions after two years. The distributions of benthic microorganisms in the Gulf can be constrained, and moreover deviations from these predictions may pinpoint impacted sites and aid in future response efforts or long-term stability studies.

## Introduction

Deep ocean sediments are the largest ecosystems on Earth and cover approximately 65% of the Earth’s surface (1). Solute transport rates in these sediments are dominated by diffusion, while larger particles are transported through burial or via bioturbation (2). The biogeochemistry regimes of deep ocean sediments are governed by the quality and quantity of deposited organic matter, most of which originated in the photic zone via photosynthesis or via transport from terrestrial sources (3,4). Furthermore, most organic matter is decomposed in the water column during transport to the seafloor (5). Therefore, oligotrophic sediments typically have very low organic carbon contents and < 20% of sedimentary organic matter is recognizable as one of the main biomolecular classes (5,6).

In contrast to the complexity and patchiness of the sedimentary carbon content, the oligotrophic deep ocean is characterized by consistent physico-chemical gradients (temperature, pressure, salinity, pH) that are often noted as the primary variables in controlling bacterial diversities (7,8). In part, this temporal and spatial stability is thought to contribute to the higher diversity and lower variation in community structure observed in deep marine sediments compared to their coastal and pelagic counterparts (3,9,10). The main environmental variables that are considered to drive microbial community composition in the deepsea are the organic carbon content, water depth, geographic distance, and overlying ecosystem productivity (9–11).

While deep marine sediments are thought to be relatively stable, they are not impervious to ecological disturbances (12–14). One glaring example of such a disturbance is the Deepwater Horizon oil spill (DWH), which represents the largest accidental marine oil spill in history, releasing an estimated 4.1–4.9 million barrels of crude oil and a third as much natural gas at approximately 1500 m water depth (15,16). Although the amounts and chemical composition of DWH oil remaining in the environment are uncertain, the largest reservoir is thought to be primarily in deep ocean sediments with a considerable, but unknown, amount deposited near coastal sediments (17). Approximately 12% of the 2 million barrels entrained in the deep ocean was transported to a 3200 km^2^ region of the seafloor (18,19). The fate of this oil will largely be determined by microbial metabolisms in these environments and by burial.

A substantial challenge in determining the impacts of this oil being deposited into deep ocean sediments is the paucity of studies on benthic microbial communities performed prior to the DWH discharge. Due to interest in petroleum exploration, previous studies in the Gulf focused almost entirely on deep subsurface sediments, or on sediments impacted by natural hydrocarbon seeps (20–22). Due to the lack of knowledge on surficial microbial communities, the majority of studies on deep ocean sediments associated with the DWH event used samples that were collected outside of impacted areas as controls (23–26). While the datasets on benthic microbes are limited to relatively few samples, results indicate that microbial communities responded quickly to oil perturbation and shifts in community composition as well as total metabolic potential were observed (23–25).

The Ixtoc-I well blowout that began in 1979 in the Gulf shares many similarities with DWH. The event released approximately 3.5 million barrels of crude oil, and represents the second largest accidental marine oil spill (27). Like DWH, Ixtoc-I crude oil was saturated with gaseous hydrocarbons and had a similar chemical composition. Approximately 25% of the total oil released is thought to have remained in seafloor sediments (27). Unfortunately, the long-term fate of Ixtoc-I oil is not well understood (28), and due to technical limitations at the time, little is known about the *in situ* microbial community response (29).

Here, we present the largest benthic marine microbial community study conducted to date, and by far the most extensive in the Gulf of Mexico. We focus on two Gulf regions that were both impacted by massive marine oil spills. Over 700 samples were analyzed representing 29 distinct sites, across four years, including archaeal and bacterial communities. Our objectives were to (1) characterize un-impacted sedimentary microbial communities to establish a baseline, (2) map the biogeographical patterns in microbial community structure across the Gulf of Mexico, (3) using this map, generate a Gulf of Mexico biogeography model of microbial community structure that can predict the abundance of dominant microbial populations, and (4) determine if impacted regions had returned to baseline conditions. We determine the environmental parameters structuring microbial communities, including the stratification of microbial communities according to sediment depth and across oxygen gradients. In addition, we address the temporal stability of microbial communities across the region.

## Material and Methods

### Sample Collection

Samples were collected during five research cruises from 2012 to 2015 across the Northern and Southern Gulf of Mexico (see Table S1, Figure 1). Sediments were sampled using a MC-800 multicorer capable of collecting up to eight, 10-cm diameter by 70 cm long cores per deployment. Cores were sectioned at either sub-centimeter (2 mm sections) or 1 cm resolution using a calibrated threaded rod attached to a core clamp stand with a tight fitting plunger (30). Sectioned sediments were frozen immediately in a portable −80 °C freezer onboard the vessel and transported back to the laboratory for DNA extraction (Table S1).

**Figure 1.**
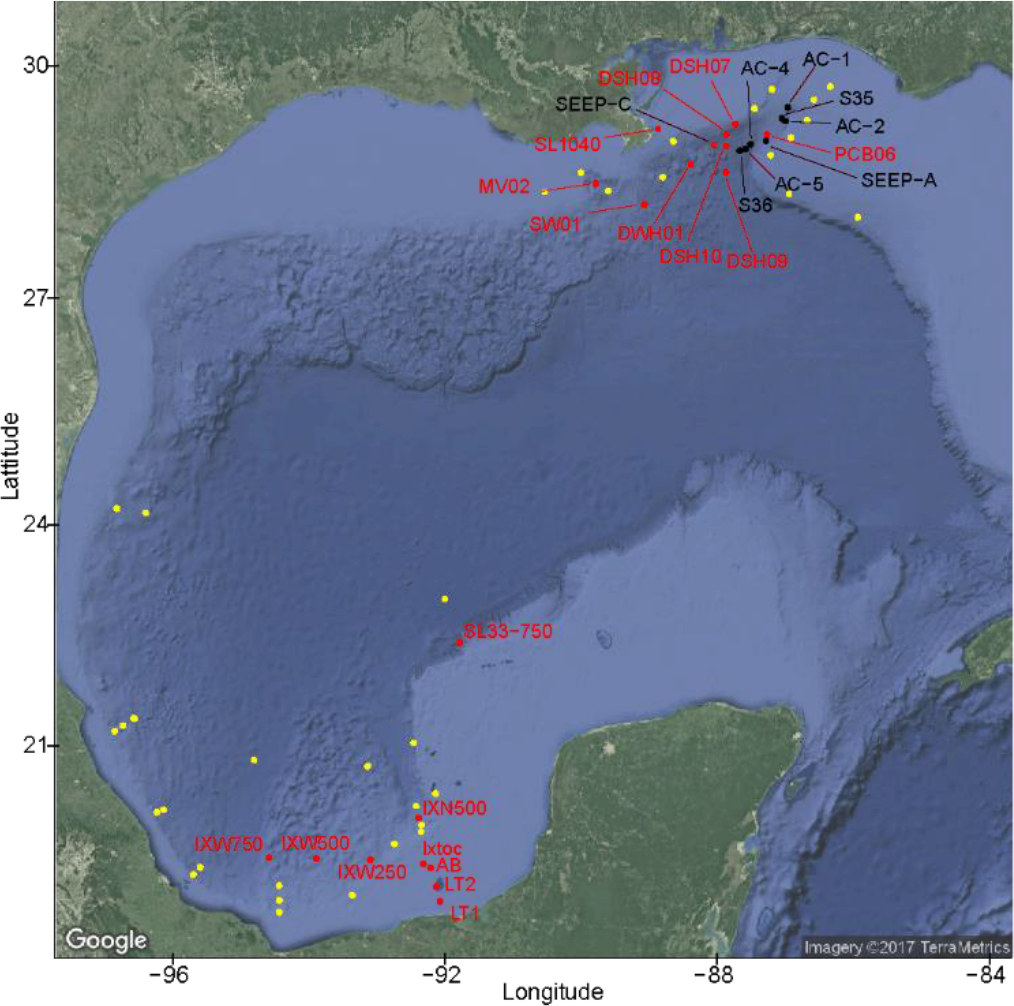
Map of the sampling sites used in this study. All yellow points are sites that were profiled for oxygen concentrations, while sites with black points are those that only have microbial community data. Sites in red are represented by both oxygen and microbial community data. All sites sampled for microbial data are labeled with the site name, which can be seen in Table S1

In the northern Gulf of Mexico (GoM), samples were collected across approximately 100,000 square kilometers, primarily focusing on the Desoto Canyon feature and near the Deepwater Horizon well-head (Table S1, Figure 1). Sampling ranged from 56 m (SL1040) near the Mississippi River Delta to 2293 m (DSH09) water depth southeast of the DWH wellhead, spanning three years (2012–2014) and four sampling time points (Table S1). In the southern Gulf of Mexico, sampling sites spanned approximately 40,000 square kilometers ranging from 16-1470 m water depth, primarily focusing on the Ixtoc-I well blowout site and along the western transect in the direction surface oil traveled (Figure 1) (31,32). Cores from the southern GoM were all collected from one cruise in 2015. For each region, sediment core profiles extended to at least 10 cm depth below the seafloor, with a subset including sections up to 20 cm of the sediment column. Our experimental design incorporated multiple levels of replication to address variation due to sampling effort, as well as spatial and temporal heterogeneity. These include: (1) technical replicates with duplicate DNA extractions performed and separate libraries generated on the same depth section, (2) multicore cast replicates where triplicate cores from the same cast were analyzed separately representing variation across approximately 1 m^2^ of the seafloor, (3) replicates at the site level where triplicate casts were deployed at the same site and cores from each cast were analyzed separately, and (4) temporal replicates where the same site was sampled up to 4 times in 3 years (Figure S1).

### Oxygen Profiles

Oxygen concentrations were profiled in the sediments using a Presens Microx4 needle-microoptode (Presens, Regensburg, Germany) along with a CHPT micromanipulator (Composite High Pressure Technologies, Lewes, Delaware, USA). Immediately after the cores were retrieved from the multicorer, a core was mounted on the extruder for stabilization and oxygen was profiled at mm-scales until the sensor measured 0 umol/L oxygen concentration. Between sites, the optode was calibrated following the manufacturer’s protocol. The depth of oxygen penetration was imported into Ocean Data View v4.6.3 (33) and predictive surfaces were generated using DIVA gridding with automatic scale lengths and with a signal to noise ratio of 50.

### DNA Extraction, SSU rRNA Gene Sequencing and Processing

Frozen sediment was sub-sectioned using a flame sterilized drillbit and total genomic DNA extractions performed on approximately 0.25 g using a MoBio Powersoil DNA extraction kit following the manufacturer’s protocol. The V4 region of the SSU rRNA was amplified using the Earth Microbiome Project 515F/806R primer set and sequenced using Illumina MiSeq 2×250 v2 chemistry (34,35). Steps are briefly outlined here and are described in detail in the Supplemental Information. Following QA/QC, operational taxonomic units (OTUs) were picked using Swarm v2.1.8 (36,37). Taxonomic affiliations for each OTU were determined using the SILVA v123 database (38). OTU counts were normalized using the metagenomeSeq package in R (39). Beta diversity was determined using Bray Curtis dissimilarity and visualized with nonmetric multidimensional scaling (NMDS) plots using the R package ‘vegan’ (40,41). Patterns in Bray Curtis dissimilarities were tested using multiple linear regressions in R (41). The most significant regression used the formula (dissimilarity ∼ sediment depth difference * water depth difference * geographic distance * Sampling Year) which was simplified to (dissimilarity ∼ sediment depth difference + water depth difference + geographic distance + sampling year + sed:water) by including only variables contributing >1% variance. Linear regression models were compared using Analysis of Variance (ANOVA), and variable contribution was determined with ANOVA (Table S2, Table S3). All plots generated use the ggplot2 package in R (41,42). All raw sequences have been uploaded to NCBI under Bioproject PRJNA414249 and all associated data and scripts can be accessed at https://doi.org/10.7266/N70G3HN3.

### Constructing a Gulf-Wide Model for Microbial Community Composition

A model for microbial community composition across the Gulf of Mexico was constructed using random forest regression implemented in the R package ‘randomForest’ at the OTU level (43). Random Forest regression is an ensemble machine learning method where multiple individual decision trees are constructed and ultimately averaged to minimize over-fitting and improve prediction accuracy. Individual decision trees are generated from a subset of the response variable (OTU abundance) and each fork or node in the tree is guided by predictor variables (randomized subset) to maximize differences in the branches. Random Forest is robust to outliers, does not assume specific distributions (nonparametric), and tests/trains the model by subsampling the response variable at each step and testing the decision node with the remaining data (11).

Water depth, sediment depth, latitude, and longitude were chosen as predictor variables and were used as proxies for sediment carbon input, oxygen concentrations, geological regime shifts from E-W and N-S as well as riverine inputs. Latitude, longitude, and water depths were extracted from the NGDC Coastal Relief model (44). The Random Forest regression model for each response variable was constructed with 500 trees, a minimum of 5 terminal nodes per tree, and randomly subsampling 1/3 of the response variable (OTU normalized counts) for each node of each tree. Only response variables of which >50% of the variance could be explained by the regression forest were retained. Details concerning the evaluation of the models are included in the Supplemental Information.

## Results and Discussion

### Environmental parameters structuring benthic microbial distributions across the Gulf of Mexico

This study incorporated sediment samples across approximately 100,000 and 40,000 square kilometers in the northern and southern Gulf of Mexico, respectively. Sampling sites ranged from 16 m to 2300 m water depth, focusing on regions adjacent to the Deepwater Horizon (DWH) and Ixtoc I blowouts. The variation in community structure across a range of spatial scales was tested at multiple replication levels (Figure S1). Technical and cast level (1 m^2^ bottom area) replicates showed similar Bray-Curtis dissimilarity values (mean = 0.23, sd= ± 0.05; 0.25 ± 0.05, respectively), while site level replicates (10-100 m^2^) were less similar (0.34 ± 0.09). When comparing the same site over time (across 3 years), lower similarity (0.54 ± 0.12) was observed, although this value may be inflated due to a small change in DNA extraction technique between 2013 and 2014 sampling years. All sites sampled at greater than 500 m water depth for the entire study show a mean dissimilarity of 0.61 ± 0.15 in comparison to a mean dissimilarity of 0.51 ± 0.13 for sites sampled at less than 100 m.

Overall, the distribution of microbial populations across these geographically distant regions of the Gulf were remarkably consistent and were structured primarily based on sediment depth along with overlying water depth (Figure 2 A, B). The sampling design focused primarily on shallow sites (<100 m) and deep sites (>1000 m), and thus the down core sediment progressions seen in Figure 2 A are structured into two main clusters primarily along the Y-axis. However, a continuous gradient with increasing water depth is expected, and indeed, is observed at the sites >1000 m water depth. Performing a multiple linear regression analysis and partitioning the variance resulted in water depth explaining 22.1% of the total variation (F = 1.29e5, p < 2e-16, Table S2).

**Figure 2.**
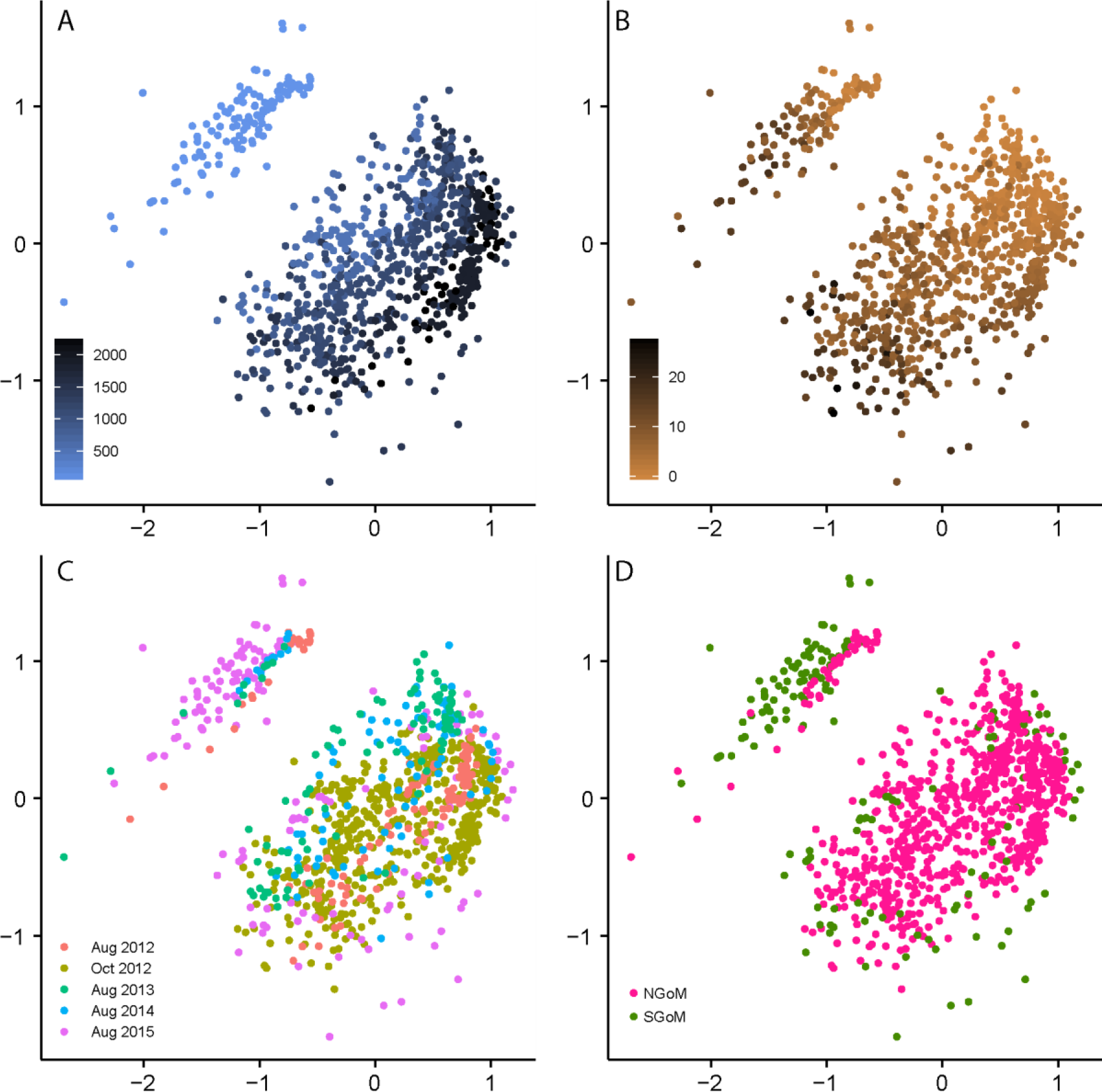
Large-scale patterns in microbial biogeography. All plots are generated from the same NMDS ordination of Bray-Curtis dissimilarity. The stress for each plot is 0.152. (A) Samples are colored based on increasing water depth. The shallow water samples (< 100m) clearly separate from the deeper water samples (> 400m). (B) Samples are colored darker with increasing sediment depths and there is a clear and consistent downcore progression from the “top-right” of the NMDS plot towards the bottom left in every core sampled. (C) Samples are colored based on collection date. (D) Cores were collected from the Southern Gulf of Mexico (colored in green) and the Northern Gulf of Mexico (colored in pink

The effect of sediment depth is primarily observed along the x-axis with surficial sediment samples clustering in the upper right of Figure 2 proceeding down core towards the bottom left, consistent with other studies (Figure 2 B) (6). Sediment depth explained 14.2% of the total microbial community variation (F = 8.3e4, p < 2e-16), and together with water depth, could explain 36% of the observed variance (74% of the total explained variance, Table S2). In contrast, geographic distance was of limited effectiveness in distinguishing microbial community compositions, explaining just 6% of the variance (F = 3.6e4, p < 2e-16), or 4% if using a discrete variable for basin region (Figure 2 C). Similarly, the sampling time point explained only 5% of the variance (F = 2.9e4, p < 2e-16) and differences were not visible in Figure 2 D. In total, water depth, sediment depth, geographic distance, sampling time point, and their interactions explained 52% of the observed variation, and a simplified model could explain 49% of the observed variance (Table S2, see Supplementary Information). Patterns in alpha diversity (Shannon, estimated OTUs, rarefaction curves) revealed lower metrics in samples collected at less than 100 m water depth compared to those collected at greater than 400 m, while changes in sediment depth were minor (Figure S2). The remarkable consistency in the Gulf seafloor microbiome may be attributed to the stability of deepsea environments, which are fueled by organic matter and nutrients supplied by the surface ocean (8). Unlike previous research, our dataset reveals that the downcore distribution of deepsea microbial communities below the surface across a large area is also likely determined by the same environmental forcings.

Previous research has mainly focused only on the biogeographic patterns of microbial communities in surface sediments, both within ocean basins and globally, and this is discussed in detail in the Supplemental Information (8,9,45–47). In general, water depth appears to have more explanatory power within a region while geographic distance and carbon bioavailability tends to have more power in global studies (1,9,10). The results generated in this study greatly contribute to the growing body of knowledge surrounding the environmental parameters structuring the seafloor microbiome. In contrast to previous work, this is the first study to examine large scale biogeographic patterns within a sediment core-profile from the shallow to the deep ocean. The larger number of samples and high depth of sequencing employed in this study along with focusing on a relatively small ocean region provided higher spatial resolution than previous work, which likely contributes to the strong biogeographic patterns in microbial community composition observed. Future global studies are encouraged to explore constraining the variation in microbial communities using a basin-based nested approach that maintains water and sediment depths as explanatory variables that facilitate the construction of predictive models for microbial communities.

### Sedimentary oxygen concentrations linked to intra-basin microbial biogeography in the Gulf of Mexico

Sedimentary oxygen profiles were determined for 42 sites across the northern and southern regions of the Gulf of Mexico using a micro-optode and micromanipulator at millimeter resolution. Oxygen profiles were obtained at most sites where cores were sectioned for microbial community composition analysis, as well as over a substantially larger area (Figure 1, Table S1). Typical core profiles are found in Figure 4 (B, D), while the interpolated maximum oxygen penetration depths across the study area are shown in Figure 3 A. At shallow, neritic sites, oxygen was depleted within the first millimeter, while oxygen penetration depths extended to 16 cm at the deepest sites (∼ 3700 m). As anticipated, oxygen penetration increased nearly linearly with increasing water depths (Figure 3 B) (48). This likely results from a greater amount of phytodetritus reaching the seafloor in shallower environments as well as increases in primary productivity closer to the coast (49,50). On the shelf, elevated chlorophyll α and β concentrations and depleted major nutrients are indicative of higher primary production (51). The oxic-anoxic interface was closer to the sediment surface in the northern Gulf in comparison to the southern Gulf at similar water depths, likely due to a lower sedimentary carbon deposition regulated by riverine inputs (Figure 3 C). The northern Gulf of Mexico is greatly influenced by the Mississippi River, the fifth largest river by volume in the world and the primary source (∼90%) of both sediment and freshwater to the entire Gulf (52–54). This region also has one of the largest and best studied anthropogenically induced “dead zones” in the world, linked to the high nutrient loadings, large phytoplankton blooms, and massive deposition events to the northern Gulf of Mexico (55,56). Furthermore, in late 2010 following the Deepwater Horizon oil spill, a massive offshore (>300 m – 1500 m sites) sedimentation event was observed whereby oily marine snow was deposited at 4-5 fold higher rates than typically observed (57,58). Relative to the northern Gulf of Mexico, the southern Gulf of Mexico generally receives lower riverine inputs (∼10% of the northern Gulf input) (54). However, our southern Gulf samples were collected in close proximity to the second largest riverine system in the Gulf of Mexico, the Grijalva-Usumacinta, as well as the smaller Coatzacoalcos and Papaloapan systems (31,59).

**Figure 3.**
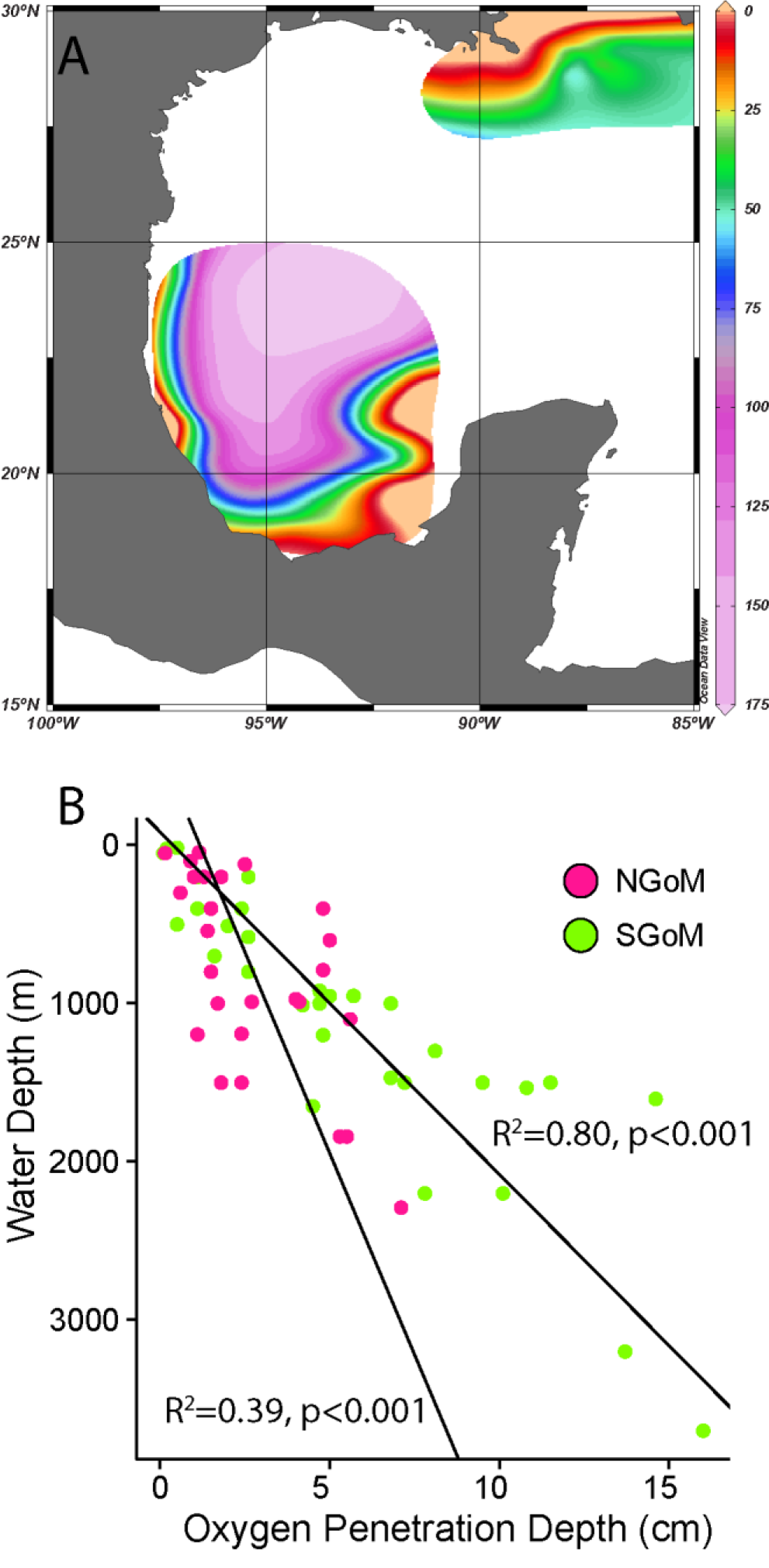
Oyxgen profiles collected throughout the Gulf of Mexico. (A) Interpolated plot showing predicted oxygen penetration depths (mm) across our study site. (B) Oxygen penetration plotted against water column depth

**Figure 4.**
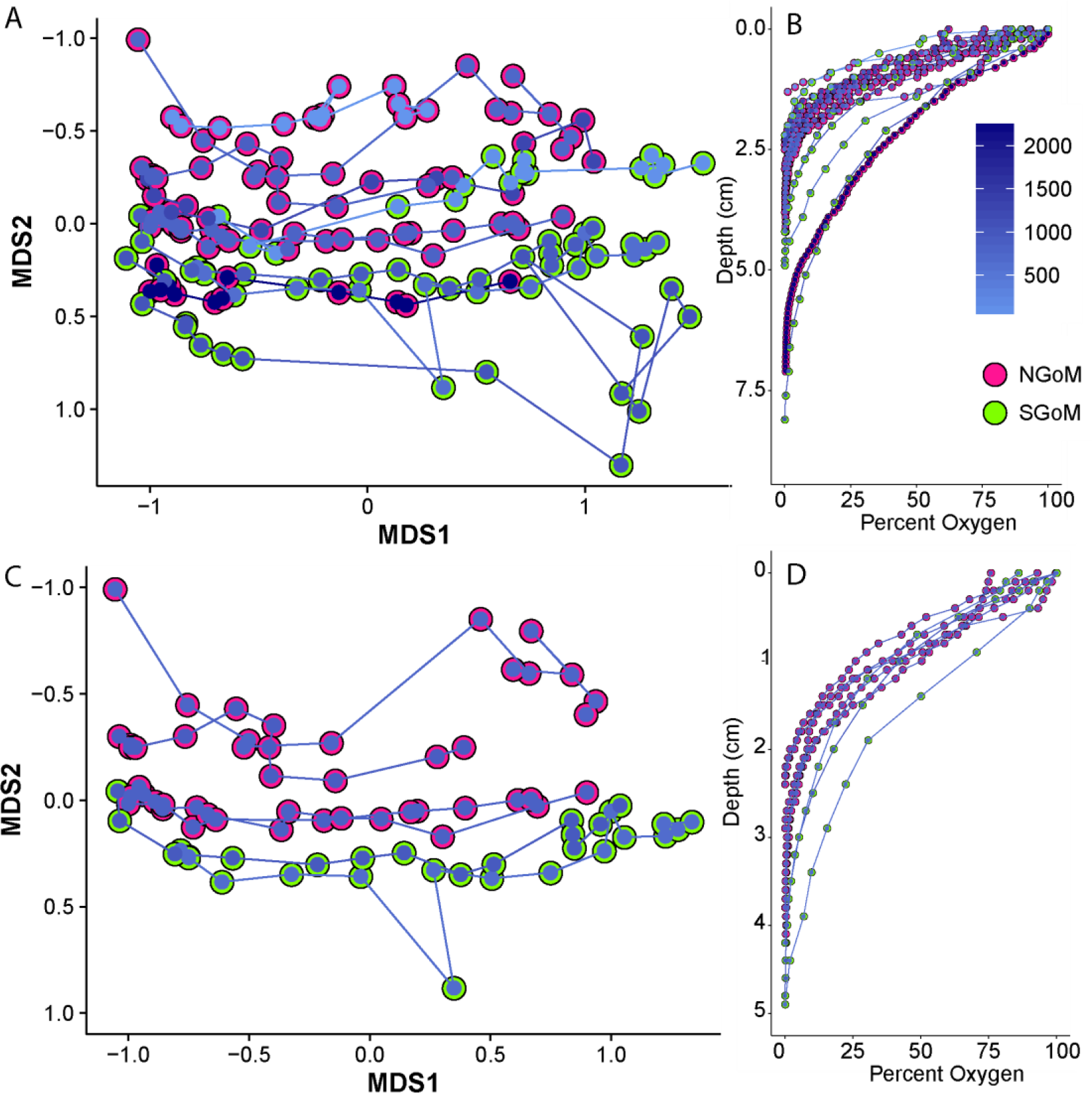
**D**ifferences in microbial community and oxygen profiles between the Northern and Southern Gulf of Mexico. The SGoM sites sampled have deeper depths of oxygen penetration and correspond to a community composition more similar to shallower sites in the NGoM. (A) NMDS of Bray-Curtis dissimilarity values for cores collected in 2014 (NGoM, pink) or 2015 (SGoM, green) from 500m – 2300m water depths. Down core progression is indicated by the connecting lines from surficial sediments on the left towards deeper samples on the right. (B) Corresponding oxygen profiles to all cores presented in A. (C, D) Same as plots A & B except only displaying sites collected between 1000-1300m.

In agreement with contrasting oxygen penetration depths (Figure 3), significant differences in microbial community composition (p < 2e-16, var. explained = 4%) were observed between the northern and southern Gulf of Mexico, although these differences were small relative to the impacts of overlying water column depth and sediment depth (Figure 2). When comparing the microbial community composition of sediments from similar water depths in each region, community composition in the northern Gulf of Mexico more closely resembled shallower sites in the southern Gulf of Mexico (Figure 4, Supplemental Information). Following a simple adjustment for water depth, the sedimentary microbial communities in these geographically distinct regions with such different quantities of riverine inputs were remarkably similar.

### The core seafloor microbiome across the Gulf of Mexico

In corroboration of previous work conducted in deep ocean oligotrophic sediments across the globe, we observed that members of the Gammaproteobacteria and Deltaproteobacteria dominate surficial communities (Figure 5, Supplemental Information). In contrast to past work, this study reveals that the MG-I Thaumarchaeota are the second most abundant class detected in the Gulf seafloor. This group likely went unreported in previous studies due to recent advances in sequencing technology and primer design (9,10). The majority of studies have focused on bacterial groups, whereas when universal primers are used, Marine Group I Thaumarchaea are shown to be very abundant (47). Although previous work in the Gulf of Mexico did not incorporate Archaea, our results on bacterial distributions are further corroborated by clone libraries of bacterial communities constructed from surficial sediments that were sampled prior to oil impacts which showed similar dominant classes (24).

**Figure 5.**
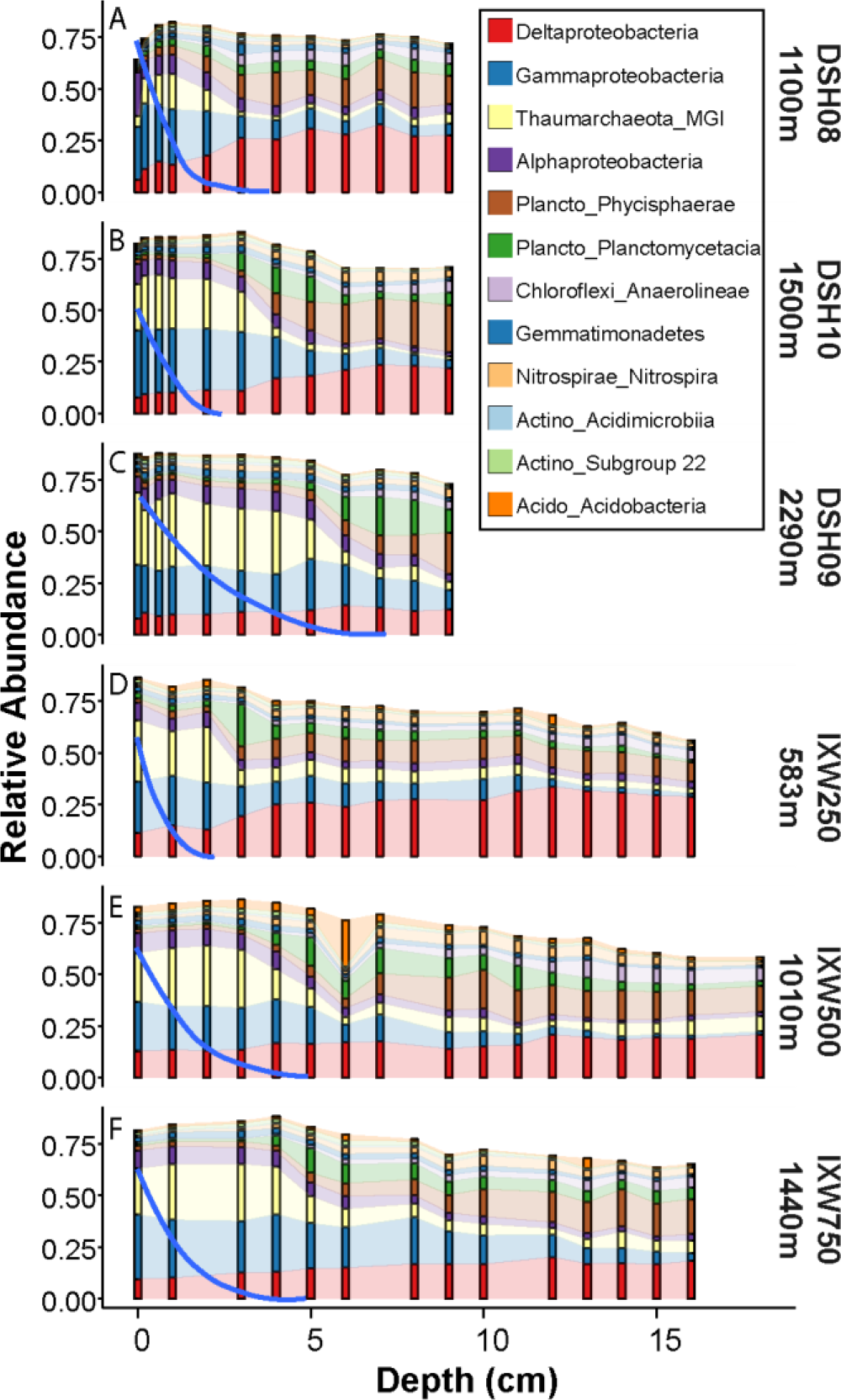
Core profiles for the 12 most abundant classes detected from deep water transects. (A-C) Representative profiles from the Northern Gulf of Mexico from the DSH core transect going offshore towards increasing water depth. (D-F) Representative profiles from the Southern Gulf of Mexico along a westward transect from the Ixtoc wellhead, also organized from shallowest to deepest site in the transect. In all plots, blue lines represent the percent oxygen concentration relative to the initial overlying water oxygen concentration.

Class level distributions along increasing sediment depths were generally consistent across core profiles at different water depths. However, a main difference was the visible inflection point in class relative abundances that coincided with oxygen depletion (Figure 5). These inflection points shifted down the sediment column with increasing water depth while becoming broader. This pattern was clearly evidenced by comparing the changes in relative abundance of the aerobic Marine Group I (Thaumarchaeota) in cores from sites DSH08 (1110 m), DSH10 (1500 m) and DSH09 (2293 m) (Figure 5). In addition, the transition between MG-I Thaumarchaeota and Phycisphaerae typically coincided with the oxygen gradient and the oxic-anoxic interface, which increased in sediment column with increasing water depth. Furthermore, this oxic-anoxic transition zone coincided with highest relative abundances of Planctomycetacia. Similar patterns, although with different methods and at different scales, have been elucidated in shallow, organic rich sediments in Aarhus Bay, where similar delineations in transition zones were linked to bioturbation activity coinciding with oxygen penetration and increased labile organic matter availability (60).

Our comprehensive dataset reveals distinct and predictable patterns in numerous OTU on a basin wide scale (Figure 6, 7c). The majority of these OTUs are not closely affiliated with characterized taxa. For example, the dataset reveals that the most abundant archaeal OTU in the Gulf seafloor microbiome is distinct from previously characterized archaea, which are mainly chemolithoautotrophic ammonia-oxidizers (Figure 6, Figure S6). The most abundant OTU within the Gammaproteobacteria is also uncharacterized and is consistently distributed at the oxic-anoxic interface (Figure S7). The most abundant and ubiquitously distributed OTU in the whole dataset is affiliated with *Syntrophobacter*, at low sequence identity (Figure S8). Given that the function of these microbial groups is not yet known, we discuss the distribution of these and many other dominant OTU further in the supplementary material. Thus, the discussion is focused on taxa putatively associated with the marine nitrogen cycle (Figure S11).

**Figure 6.**
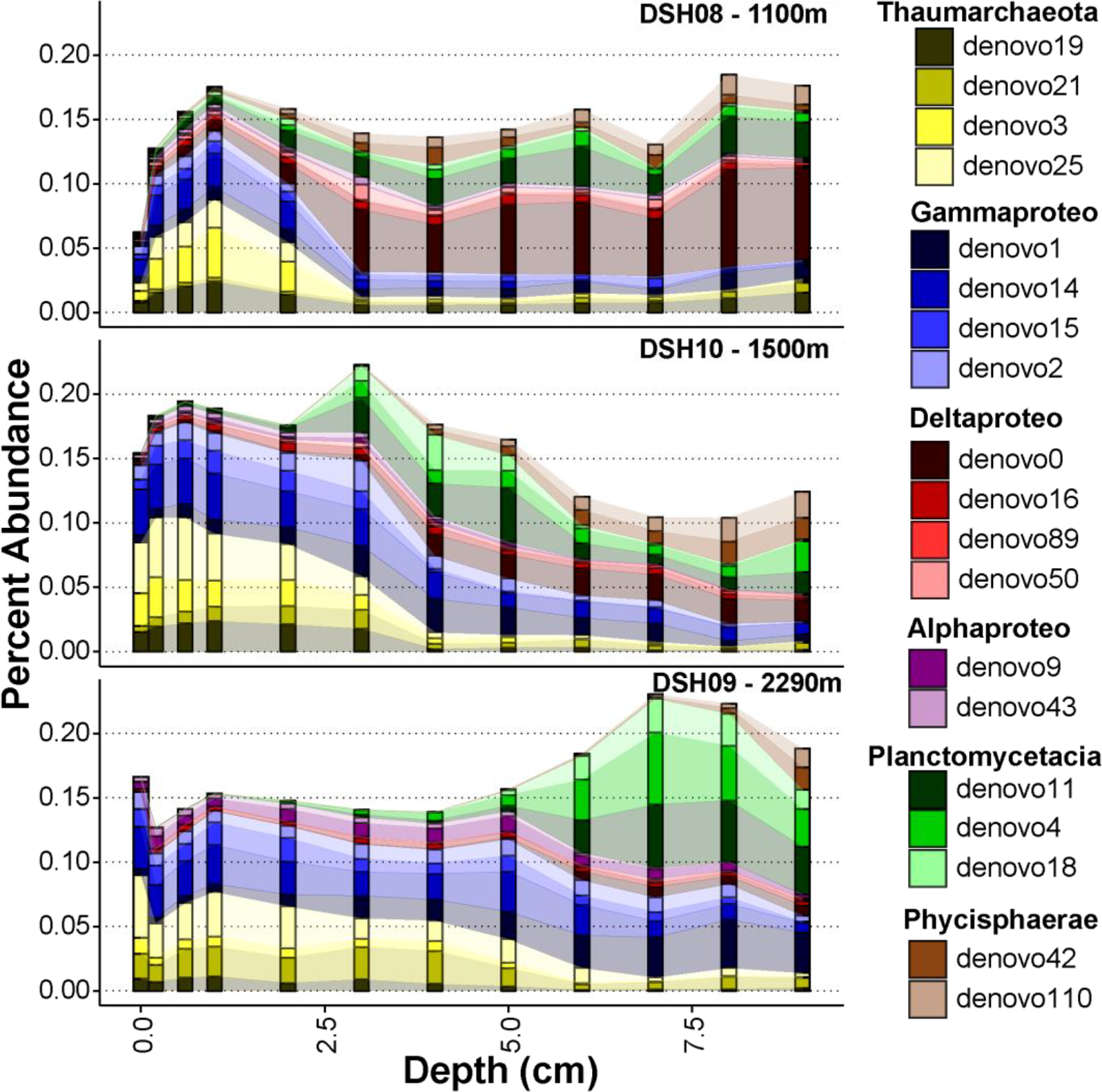
Core profiles representing deep water sites in the Northern Gulf of Mexico showing population (OTU) level distributions with increasing sediment and water depths. Core profiles are taken from the DSH transect and are arranged in order of increasing water depth. See Figure S4 for similar trends within the Southern Gulf of Mexico.

The four most abundant Marine Group I OTU (Thaumarchaeota) were all phylogenetically related to *Nitrosopumilus maritimus* (3 OTU >97% sequence identity and 1 OTU at 93% identity). Distributions of these four abundant *Nitrosopumilus-*like OTU were not driven by depth within the sediment column as was found by Durbin and Teske, and instead were related to the overlying water depth, showing putative niche differences due to organic matter inputs (2010) (Figure S6). Marine Group I Thaumarchaeota are well known members of oxic and suboxic sediments (1,47,61–65). All characterized isolates similar to these 4 OTU are aerobic autotrophs capable of oxidizing ammonia at very low concentrations. Increases in their relative abundances have been shown to be concurrent with decreases in the quality and quantity of organic carbon within sediments (47). It has also been shown that in deep, low energy pelagic environments, a large portion of *Nitrosopumilus*-like populations were capable of using urea as a carbon and ammonia source to fuel autotrophic nitrification (66).

All dominant OTU affiliated with the Planctomycetacia showed high sequence identity to *Candidatus Scalindua* spp (99-100%). These dominant OTU are expected to be anaerobic ammonium oxidizing bacteria (anammox), which are chemoautotrophic, ubiquitously found in anoxic systems, and couple the oxidation of ammonium with nitrite as the electron acceptor (67,68). Members of this genus tend to be psychrophilic, and are inhibited by low sulfide concentrations (4-100 µM), which potentially explains their depth limits in our study, although it may also be due to ammonium or nitrite availability (67). The Planctomycetes were nearly absent in the shallowest sites, and absent in surficial sediments; in agreement with previous studies that indicate ca. *Scalindua* is more abundant in oligotrophic systems (47,69). All dominant Planctomycetes OTU tended to reach a maximal relative abundance at mid-sediment depths below the oxic-anoxic interface with an overlapping distribution to the *Nitrosopumulis*-like OTUs (Figures 6, S4, S10). Due to their overlapping distributions, ammonium oxidizing archaea may provide a nitrite source for ca. *Scalindua* (70).

In parallel with the distribution of ca. *Scalindua*-like microbial populations, putative bacterial ammonium oxidizers (*Nitrosococcus-like*, and an unclassified Nitrosomonadaceae OTU) were also absent in nearshore sediments or in surficial sediments from deep ocean sites and exhibited an increase in relative abundance with increasing water depth (Figure S11). Abundant putative nitrite oxidizers affiliated with *Nitrospira* and *Nitropina* groups were detected that were also only present in deep water sediments below the aerobic-anaerobic transition zone, similar to the *Scalindua-*like population distributions. While circumstantial, the relatively high abundances of these putative groups in deep ocean sediments provides evidence of inorganic nitrogen species being important electron acceptors and donors in deep Gulf of Mexico sediments, particularly for fueling autotrophic metabolisms. These observations are supported by previous studies showing the complexity of nitrogen cycling in similar sediments as well as the growing body of knowledge towards the importance of chemolithoautotrophs in oligotrophic deep ocean sediments (1,71–73)

### Construction of a predictive microbial community composition model throughout the Gulf of Mexico

After visualizing the consistent biogeographic patterns portrayed in Figure 2, it was recognized that a regression-based model may effectively reconstruct the relative abundances of microbial groups across these regions. A machine learning method, random forest regression, was employed that has previously been successful in population distribution models (11,74–79). As stated in the methods section, the model was constructed using the variables water depth, sediment depth, latitude, and longitude as proxies for carbon input, redox state, and geographic variations due to riverine inputs and differing geochronology across the Gulf of Mexico (57).

This analysis was run for every abundantly detected OTU (9588 OTU). All OTU with greater than 50% variance explained by the full model were retained (4129 OTU). Overall, the models fit the observed data well (R^2^ = 0.88). Residuals were weakly correlated with OTU abundance as the model tends to underestimate the very abundant OTU. The model was further tested by calculating the root mean squared error (RMSE) compared to 1000 bootstrapped randomly permuted RMSE and was found to be highly significant (p < 2e^-16^, Z-test).

To verify the utility of the model, model predictions were compared with observed counts for the dominant OTU (Figure S5, S6-S10). We anticipate that the model can be used as a tool to assess future impacted sediments that may be poorly characterized and to test this, we hindcasted the microbial community composition at previously impacted sediments. This was accomplished by calculating the total root mean squared error (RMSE) for each sample using observed OTU abundance and predicted OTU abundances, including the Mason et al. (2014) surficial sample set (Figure 7 C). The higher the RMSE value for each sample, the great the observed community composition deviated from the model prediction. The most impacted sites had higher RMSE values than the “below EPA limits” sites, although both were skewed to the right of the model predictions for our data (Figure 7 C). Therefore, while controlling for experimental based variation, this model can still be effective at determining impacts.

**Figure 7.**
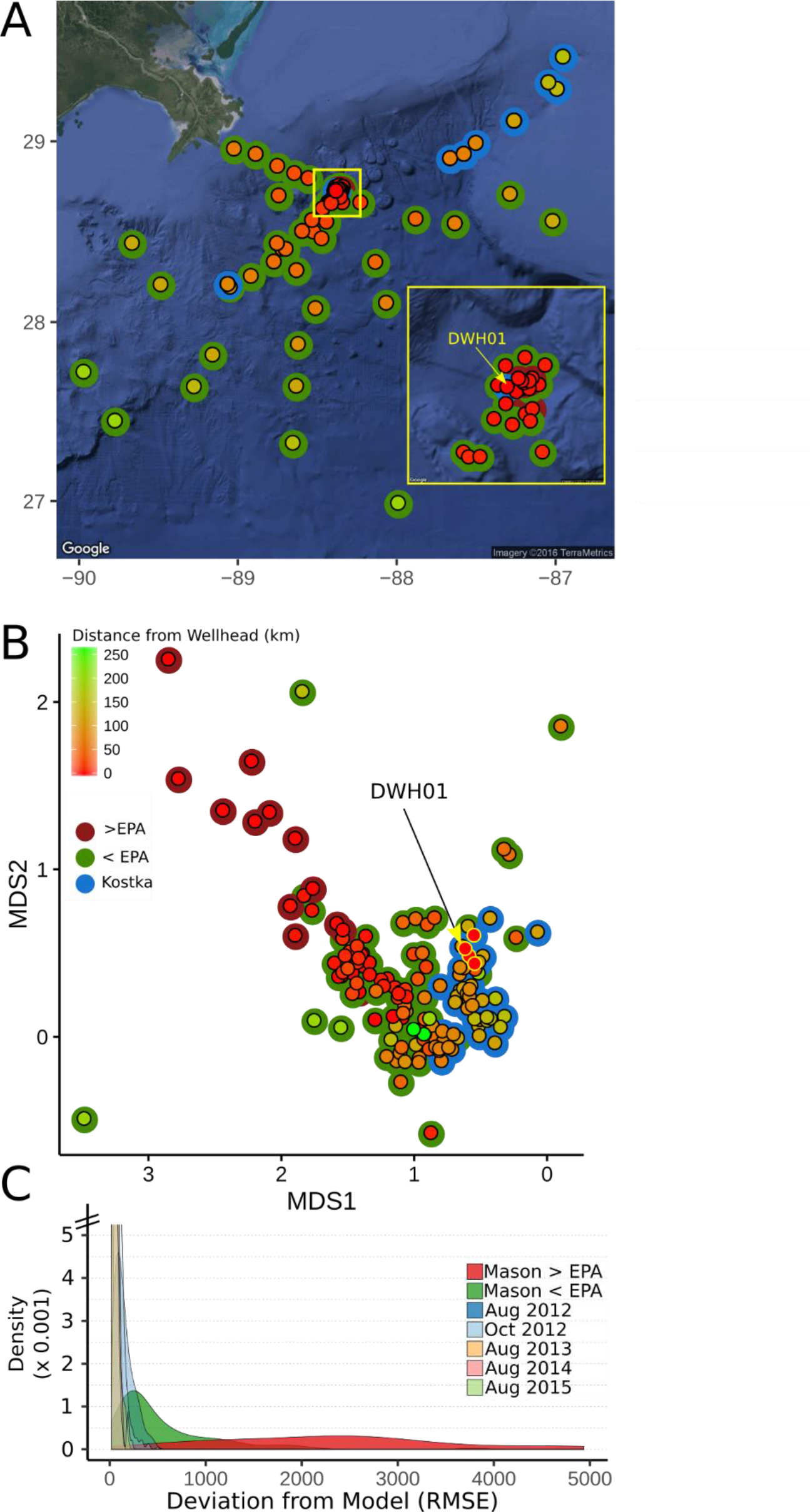
Meta-study revealing localized impact and subsequent recovery. Data generated by Mason et al., 2014 from surficial samples (red, green) was re-analyzed with surficial samples from this study (blue). (A) Geographic map plotting each sampling location. Samples outlined in red from Mason et al., exceeded EPA benchmarks while samples outlined in green were below EPA benchmarks. The fill color for each point represents distance from well head, going from red to green with increasing distance. (B) NMDS of Bray-Curtis dissimilarity values, plotted using the same color scheme as in A. The wellhead samples (DWH01) are indicated by yellow outlines and were collected in 2012 and 2013. (C) Deviations of every sample collected, including the full sediment core for samples from this study, from the OTU distribution models. Samples are colored by the sampling date or by oil contamination state. High RMSE (root mean squared error) values represent samples that strongly differ from OTU relative abundance predictions.

### Impacts of the DWH and internal testing of the predictive model

The earliest samples included in this study were collected in August 2012, over two years after the Deepwater Horizon well blowout. No evidence of a large oil spill signature was detected in sediments sampled from areas known to be impacted by sedimentary oil deposition related to the DWH blowout (18,80–82). At the broad biogeographic scale visualized in Figure 2, the lack of an “oil-impacted cluster” indicated that any potential impact on community composition was small relative to the strong environmental gradients influenced primarily by water depth and sediment depth. To compare our data explicitly with sediments that were shown to be strongly impacted by oil, a meta-analysis based approach was employed. We observe that all surficial sediment samples from this study, including those collected close to the DWH wellhead, clustered adjacent to samples classified as below EPA limits and were distinct from those classified as most impacted by oiling (23) (Figure 7 B). Furthermore, the similarities between the below EPA limit samples and our dataset indicated a temporally stable and Gulf-wide conserved microbial community relative to a strong pulse disturbance. Finally, previously impacted sediments that were collected and analyzed during this study did not deviate from model predictions, unlike those collected in 2010 (Figure 7 C).

These results are corroborated by a previous study that detailed a microbial community succession event in oil impacted surficial sediments immediately following the oiled marine snow sedimentation event, through the generation of full length SSU rRNA gene clone libraries (24). The authors suggested that by November 2010 the microbial community composition once again resembled pre-impacted surficial microbial communities (24). Our first samples were collected in August 2012 and much of our dataset comes from 2013 and 2014 for the NGoM, years following a potential return to baseline condition. Over much larger scales, our results do not show a disturbance signature in cores collected at the Ixtoc-I wellhead or within the PEMEX (Mexican state-owned petroleum company) restricted area that has extensive oil extraction pressures. The lack of an Ixtoc-I signature in the microbial community is expected considering the resilience observed in the sedimentary communities in the northern Gulf of Mexico, and the lack of a PEMEX oil signature indicates we did not sample sediments that were recently exposed to oil.

## Conclusions

The primary objectives of this study were to define the core microbiome across the Gulf of Mexico seafloor, characterize biogeographic patterns in microbial populations in Gulf sediments, and use these results to constrain the impacts of petroleum hydrocarbons from major oil spills to sedimentary microbial communities. Using the largest marine sediment microbial community composition dataset generated to date, we show that the distribution of microbial communities is broadly consistent across the entire region in terms of the OTU detected and their relative abundances. We further demonstrate that the distribution of seafloor microbial communities is well approximated by the overlying water depth and depth within the sediment column, which together explain 38% of our observed variation, regardless of geographic sampling region. The small differences between the regions are potentially related to the amount of organic carbon reaching the seafloor, with the southern Gulf showing greater depths of oxygen penetration as well as microbial communities that were more similar to deeper-water counterparts in the northern Gulf. Microbial community composition across the Gulf revealed similar taxonomic groups as observed in other marine sediment biogeography studies across the globe and is dominated primarily by groups that are poorly characterized with unknown ecological functions within the sediment. Putative ammonium oxidizing archaea (*Nitrosopumilus*-like OTU) were among the most relatively abundant groups detected in the dataset across oxygenated sediments in the Gulf of Mexico, while other putative chemolithoautotrophic nitrogen-transforming groups were relatively abundant only in the deep ocean sediments. Overall, the dominant OTU detected from each of the major classes, generally had low sequence identities to characterized taxa.

The lack of a strongly perturbed community in our dataset, along with the incorporation of previous studies demonstrating an oil related perturbation signal, suggests that impacted sediment communities had returned or are returning to baseline conditions. Finally, the biogeographical distributions of seafloor microbial populations were used to generate a predictive model for over 4000 dominant populations that relies on easy-to-obtain geospatial variables. With this model, it is possible to predict community compositions across the Gulf of Mexico, and use deviations from such a prediction as a tool to assess future impacted sediments that may be poorly characterized along with natural ecosystems that are conspicuously different from oligotrophic ecosystems such as cold seeps.

## Supporting information

Full Supplementary Information

## Acknowledgements

This work was made possible in part by a grant from The Gulf of Mexico Research Initiative to the C- IMAGE II consortium and in part by a National Science Foundation Graduate Research Fellowship (WAO) under grant no. 2013172310. The authors would like to thank the ship crews on the R/V Weatherbird III and the R/V Justo Sierra that made the sample collection possible. Data are publicly available through the Gulf of Mexico Research Initiative Information and Data Cooperative (GRIIDC) at https://data.gulfresearchinitiative.org (doi: 10.7266/N70G3HN3). Any opinions, findings, and conclusions or recommendations expressed in this material are those of the authors and do not necessarily reflect the views of the National Science Foundation.

