## Supplementary material for "Defining the seafloor microbiome of the Gulf of Mexico and its response to oil perturbation": Full Supplementary Information

Running title: Deepsea benthic microbial biogeography

### Methods

#### *SSU rRNA Gene Sequencing*

All samples from 2012-2013 were generated by the Michigan State University sequencing facility. Raw DNA extracts were transferred frozen and were amplified with barcoded 515F/806R primers according the Earth Microbiome Project standard protocol (Caporaso et al., 2012; http://www.earthmicrobiome.org/emp-standard-protocols/16s/). A total of 5 MiSeq 2x250 bp runs were performed and demultiplexed sequences were transferred to Georgia Institute of Technology. Samples collected from 2014-2015 were processed in house using the same primer set targeting the V4 region, however, we employed the Fluidigm Access Array kit to barcode the samples post-PCR amplification. Multiplexed samples were pooled at equimolar ratios and paired-end sequenced (2x250 bp) on an Illumina MiSeq following the protocol outlined in by Green and colleagues (Green *et al.*, 2015). In brief, the 515F/806R primer set was modified with linker sequences at the 5' ends of each primer and the V4 SSU rRNA gene region was amplified. A second stage PCR was performed using a primer set that contains the Illumina sequencing adapters along with sample-specific barcodes (multiplexing stage). At this stage, sample amplicons were pooled at an equimolar ratio and were ready for sequencing.

#### *Calculating Alpha Diversity Metrics*

Alpha diversity metrics were calculated using the normalized OTU table as described in the methods section. Rarefaction curves were generated using a wrapper function created by B. Hausmann and described here (https://github.com/joey711/phyloseq/issues/143). Shannon indices (base e) were determined using the R package 'vegan' and the estimated richness was calculated by fitting a zero-truncated negative binomial model by the R package 'preseqR' (Deng et al., 2014).

### Results

#### Populations Specific Niche Differences

##### *MG-I Archaea*

Using the full dataset, the most abundant MG-I OTU (denovo3, 93% identity to *N. maritimus*) dominated sites sampled at approximately 1000 m water depth (Figures 6, Figure S6), while denovo25 (98% to *N. maritimus*) was abundant in the deepest sites. Conversely, denovo19 and denovo21 (97%, 99% to *N. maritimus* respectively) were most abundant at the shallowest sites. However, all four were abundant in surficial sediments across the entire Gulf and did not have a strong differential response to sediment column depth as was found by Durbin and Teske (2010). Instead of oxygen concentrations dictating the niches of these specific OTU (e.g. specific adaptation to low oxygen concentrations), it may be organic matter quantity and quality that structure the niches, as has been shown in surficial sediments in Antarctica (Learman *et al.*, 2016). While water depth best explains these patterns, which governs organic matter inputs in the Gulf, there were latitudinal and longitudinal differences in the dominant MG-I OTU (Figures S3, S6). However, these smaller shifts due to spatial variation were also likely driven by organic matter quality and quantity, which vary based on proximity to the Mississippi river delta across the Gulf (Goñi et al., 1997).

##### *Gammaproteobacteria*

Our most abundant Gammaproteobacteria OTU (denovo1) was 96% similar to the only isolated strain, *Woeseia oceani*, and showed maximum relative abundances just below the oxic-anoxic interface and in deeper water sites, similar to members of the Planctomycetacia group (Figure 6, Figure S7). *Woeseia*-like population may be respiring nitrate or nitrite coupled with chemoorganoheterotrophy (Dyksma *et al.*, 2016; Mußmann *et al.*, 2017). Unlike denovo1, the remaining dominant

Gammaproteobacterial OTU were most abundant in surficial, aerobic sediments of the deep ocean, although their abundances also increased with increasing water depth (Figure 6, S4, S7). Denovo14 and denovo15 were both assigned to the JTB255/*Woeseiaceae* group (95% and 98% similarity to denovo1 respectively). The second most abundant OTU, denovo2, was divergent from the JTB255 group and showed 93% sequence identity to denovo1. The closest BLAST hit was to the isolated strain *Thioprofundum hispidum*, at 94% sequence identity. It was < 95% sequence identity to denovo14 and denovo15, although they exhibit similar distribution patterns. While the phylogenetic identity of denovo2 was less certain, *T. hispidum* was characterized as a chemolithoautotrophic sulfur-oxidizing strain within the Chromatiales order (Mori *et al.*, 2011). With the observed distribution patterns, these 3 OTU (denovo14, 15, 2) were likely utilizing oxygen as a terminal electron acceptor either for chemolithoautotrophic processes observed within *Thioprofundum spp.* coupled with sulfur oxidation or as aerobic chemoorganotrophs as observed in the JTB255/*Woeseiaceae* group (Mori *et al.*, 2011; Mußmann *et al.*, 2017).

##### *Deltaproteobacteria*

Similar to the Gammaproteobacteria, little overlap was observed between Deltaproteobacterial OTU associated with the shallower sites and those in the deeper sites. In deep ocean sediments, most of the sequences affiliated with Deltaproteobacteria, belong to denovo0. This was the most abundant OTU detected in our dataset, and its relative abundance increased with increasing sediment depth. Interestingly, maximum relative abundance of denovo0 was found at intermediate water column depths (1000 – 1200 m) (Figure 6, S8). Although not closely affiliated to any cultivated strain, denovo0 was most similar to *Syntrophobacter fumaroxidans* at 91% and assigned to the *Syntrophobacteraceae* family by SILVA. The family consists of strictly anaerobic fermentative or sulfate respiring members that are broadly distributed (Kuever, 2014). The other dominant deep ocean sediment Deltaproteobacterial OTU, denovo16 (<90% similarity to denvo0), showed similar a distribution to denovo0, and did not decrease with increasing water depth. Denovo16 showed 91% sequence similarity to *Deferrisoma camini*, a thermophilic strictly anaerobic iron-reducing bacterium, within the uncharacterized and uncultivated NB1-J group. In the shallow sites, denovo89 and denovo50 were very abundant throughout the sediment column (Figure S8). Denovo50 shares 96% identity to *Desulfatiglans aniline*, and was more abundant in the shallow SGoM than the shallow NGoM, and was not abundant at deeper sites. Of note, denovo89 was closely affiliated (98% identity) with the deep ocean OTU denovo0, and thus also likely represents a poorly characterized population within the *Syntrophobacteraceae* family.

*Planctomycetacia*

All dominant OTU affiliated with the Planctomycetacia were closely related to *Candidatus Scalindua* spp (99-100%). These dominant OTU are likely anaerobic ammonium oxidizing bacteria (anammox), which are chemoautotrophic, ubiquitously found in anoxic systems, and couple the oxidation of ammonium with nitrite as the electron acceptor (Penton *et al.*, 2006; Oshiki *et al.*, 2016). Members from this genus tend to be psychrophilic, and are inhibited by low sulfide concentrations (4-100 µM), which potentially explains their depth limits in our study, although it may also be due to ammonium or nitrite availability (Oshiki *et al.*, 2016). The Planctomycetes were nearly absent in the shallowest sites, and absent in surficial sediments, in agreement with previous studies that indicate ca. *Scalindua* is more abundant in oligotrophic systems (Canion *et al.*, 2014; Learman *et al.*, 2016). All dominant Planctomycetes OTU tended to reach a maximal relative abundance at mid-sediment depths below the oxic-anoxic interface with an overlapping distribution to the *Nitrosopumulis*-like OTUs (Figures 6, S4, S9). Ammonium oxidizing archaea may provide a nitrite source *Scalindua*, and the mostly exclusive zonation patterns observed may limit direct competition for ammonium (Lipsewers *et al.*, 2014). In some sites, a clear “bubble” in abundance surrounded this zone, while in other sites they persist through the top 10-20 cm (Figures 6, S4, S9). Denovo4 tended to be more abundant with increasing water depth, while denovo11 had higher relative abundances in the 600-1200 m water depth range. Denovo18 was co-localized with denovo4 although this population tended to be deeper in the sediment profile (Figure 6). Previous studies focusing on surficial sediments typically found low Planctomycetacia abundances (Schauer *et al.*, 2010; Bienhold *et al.*, 2016; Learman *et al.*, 2016) while studies that included a larger depth range, or core profiles found higher abundances with depth than at the surface (Zinger *et al.*, 2011; Canion *et al.*, 2014).

*Phycisphaerae*

The dominant Phycisphaerae OTU were only abundant in the deep ocean and different OTU were present in the shallower sites (Figure S10). All increased in relative abundance with increasing sediment depth and were only present below the aerobic to anaerobic transition zone. These OTU were more relatively abundant in the NGoM than the SGoM although the same OTUs were detected in both regions (Figures 6, S4, S10). The more abundant OTU, denovo42, was affiliated with an uncharacterized family, MSBL9, in SILVA taxonomy with no closely related named strains (< 80% sequence identity) although closest BLAST hits were from anaerobic marine sediments with a 100% 16S clone sequence from Gulf of Mexico sediments (Reed *et al.*, 2006). Denovo110 was also affiliated with and uncharacterized family, MLA10 with close BLAST hits from clone sequences generated from anoxic marine sediments.

**Table S1.** List of all sites used in this study.

| Sampling Site | Water Depth (m) | Latitude | Longitude | Number of Unique Samplings | Number of Years | 16S Sequences | Oxygen Profile |
|---|---|---|---|---|---|---|---|
| Abkatum | 50 | 19.314270 | -92.208000 | 1 | 1 | Y | Y |
| AC-1 | 490 | 29.474500 | -86.958700 | 1 | 1 | Y | Y |
| AC-2 | 845 | 29.297600 | -86.996800 | 1 | 1 | Y | Y |
| AC-4 | 1732 | 29.000300 | -87.507400 | 1 | 1 | Y | Y |
| AC-5 | 1850 | 28.940100 | -87.582400 | 1 | 1 | Y | Y |
| DSH07 | 400 | 29.255817 | -87.732383 | 1 | 1 | Y | Y |
| DSH08 | 1100 | 29.122783 | -87.867733 | 1 | 1 | Y | Y |
| DSH08 | 1100 | 29.122783 | -87.867733 | 3 | 3 | Y | Y |
| DSH09 | 2290 | 28.636500 | -87.868500 | 2 | 2 | Y | Y |
| DSH10 | 1500 | 28.979050 | -87.891617 | 4 | 4 | Y | Y |
| DWH01 | 1500 | 28.736669 | -88.387161 | 2 | 2 | Y | Y |
| IWX250 | 583 | 19.430680 | -93.094970 | 1 | 1 | Y | Y |
| IXN500 | 1200 | 20.014000 | -92.389000 | 1 | 1 | Y | Y |
| IXTOC | 60 | 19.370080 | -92.317180 | 2 | 1 | Y | Y |
| IXW500 | 1010 | 19.443000 | -93.889000 | 1 | 1 | Y | Y |
| IXW750 | 1470 | 19.458000 | -94.587000 | 1 | 1 | Y | Y |
| LT1 | 16 | 18.808000 | -92.057000 | 1 | 1 | Y | Y |
| LT2 | 21 | 19.061000 | -92.127000 | 1 | 1 | Y | Y |
| MV02 | 541 | 28.494167 | -89.779367 | 2 | 2 | Y | Y |
| PCB06 | 990 | 29.122700 | -87.266217 | 3 | 3 | Y | Y |
| PCB06 | 990 | 29.122700 | -87.266217 | 1 | 1 | Y | Y |
| S35 | 670 | 29.333600 | -87.050200 | 1 | 1 | Y | Y |
| S36 | 1841 | 28.916400 | -87.669100 | 1 | 1 | Y | Y |
| SE02 | 972 | 28.359167 | -86.944267 | 1 | 1 | Y | Y |
| SEEP-A | 1110 | 29.043000 | -87.282400 | 1 | 1 | Y | No |
| SEEP-C | 1116 | 28.990000 | -88.045500 | 1 | 1 | Y | No |
| SL1040 | 50 | 29.196050 | -88.868833 | 3 | 3 | Y | Y |
| SL33-750 | 1300 | 22.415000 | -91.785000 | 1 | 1 | Y | Y |
| SW01 | 1192 | 28.220867 | -89.069500 | 4 | 4 | Y | Y |
| SL26A250 | 500 | 21.199 | -96.851 | 1 | 1 | No | Y |
| SL26A750 | 1500 | 21.37 | -96.559 | 1 | 1 | No | Y |
| IXNW1600 | 3200 | 20.804 | -94.803 | 1 | 1 | No | Y |
| 23-92 | 3700 | 23 | -92 | 1 | 1 | No | Y |
| SL33-200 | 400 | 22.333 | -91.7 | 1 | 1 | No | Y |
| SL31A-1000 | 2200 | 20.718 | -93.137 | 1 | 1 | No | Y |
| IXN1000 | 2200 | 20.345 | -92.137 | 1 | 1 | No | Y |

| | | | | | | | |
|---|---|---|---|---|---|---|---|
| IXN750 | 1650 | 20.172 | -92.423 | 1 | 1 | No | Y |
| IXN250 | 800 | 19.912 | -92.342 | 1 | 1 | No | Y |
| IXN100 | 400 | 19.817 | -92.349 | 1 | 1 | No | Y |
| SL31-100 | 200 | 21.038 | -92.46 | 1 | 1 | No | Y |
| Ixtoc 1 | 50 | 19.366 | -92.313 | 1 | 1 | No | Y |
| Abkatun | 50 | 19.913 | -92.206 | 1 | 1 | No | Y |
| LT3 | 50 | 19.36 | -92.28 | 1 | 1 | No | Y |
| IXNW350 | 700 | 19.645 | -92.741 | 1 | 1 | No | Y |
| IXW250 | 580 | 19.429 | -93.094 | 1 | 1 | No | Y |
| SL30A-100 | 200 | 18.936 | -93.36 | 1 | 1 | No | Y |
| SL30-500 | 920 | 19.071 | -94.433 | 1 | 1 | No | Y |
| SL30-250 | 510 | 18.866 | -94.433 | 1 | 1 | No | Y |
| SL30-100 | 200 | 18.7 | -94.433 | 1 | 1 | No | Y |
| SL28-750 | 1500 | 19.324 | -95.594 | 1 | 1 | No | Y |
| SL28-500 | 1000 | 19.224 | -95.696 | 1 | 1 | No | Y |
| SL27-750 | 1500 | 20.12 | -96.133 | 1 | 1 | No | Y |
| SL27-500 | 1000 | 20.085 | -96.233 | 1 | 1 | No | Y |
| SL26-750 | 1533 | 21.37648 | -96.57455 | 1 | 1 | No | Y |
| SL26-500 | 953 | 21.27383 | -96.72933 | 1 | 1 | No | Y |
| SL25-750 | 1603 | 24.15995 | -96.39425 | 1 | 1 | No | Y |
| SL25-500 | 952 | 24.21741 | -96.82131 | 1 | 1 | No | Y |
| PCB06 | 990 | 29.1227 | -87.26622 | 1 | 1 | No | Y |
| PCB03 | 100 | 29.73833 | -86.33833 | 1 | 1 | No | Y |
| SL7150 | 200 | 29.56833 | -86.57833 | 1 | 1 | No | Y |
| MC04 | 400 | 29.30575 | -86.67637 | 1 | 1 | No | Y |
| DSH08 | 1100 | 29.12278 | -87.86773 | 1 | 1 | No | Y |
| DSH07 | 400 | 29.25582 | -87.73238 | 1 | 1 | No | Y |
| SL1040 | 50 | 29.19605 | -88.86883 | 1 | 1 | No | Y |
| DWH01 | 1500 | 28.73667 | -88.38716 | 1 | 1 | No | Y |
| HC01 | 45 | 28.38413 | -90.53097 | 1 | 1 | No | Y |
| MV02 | 541 | 28.49417 | -89.77937 | 1 | 1 | No | Y |
| SW01 | 1192 | 28.22087 | -89.0695 | 1 | 1 | No | Y |
| SW03 | 1196 | 28.57558 | -88.80043 | 1 | 1 | No | Y |
| SE02 | 972 | 28.35917 | -86.94427 | 1 | 1 | No | Y |
| DSH09 | 2290 | 28.6365 | -87.8685 | 1 | 1 | No | Y |
| S36 | 1841 | 28.9164 | -87.6691 | 1 | 1 | No | Y |
| NT800 | 789 | 28.056 | -85.93351 | 1 | 1 | No | Y |
| MC06 | 600 | 29.08355 | -86.91452 | 1 | 1 | No | Y |
| SL8100 | 200 | 29.70167 | -87.19167 | 1 | 1 | No | Y |
| SL1460 | 120 | 29.45648 | -87.45088 | 1 | 1 | No | Y |
| DSH10 | 1500 | 28.97905 | -87.89162 | 1 | 1 | No | Y |

| | | | | | | | |
|---|---|---|---|---|---|---|---|
| PCB09 | 1000 | 28.85912 | -87.21468 | 1 | 1 | No | Y |
| SL11150 | 300 | 29.03785 | -88.64423 | 1 | 1 | No | Y |
| MV03 | 800 | 28.39825 | -89.59938 | 1 | 1 | No | Y |
| SL16150 | 200 | 28.63537 | -90.0015 | 1 | 1 | No | Y |

Table S2. Analysis of variance table for multiple linear regressions used in this study.

Simplified best fit model.

| | Df | Sum Sq | Mean Sq | F value | Pr(>F) | Percent Variance Explained |
|---|---|---|---|---|---|---|
| water_dif | 1 | 1475.448 | 1.48E+03 | 129176.4 | 0 | 22.09 |
| sed_dif | 1 | 951.5594 | 9.52E+02 | 83309.6 | 0 | 14.25 |
| geo_dist | 1 | 405.9833 | 4.06E+02 | 35544.09 | 0 | 6.08 |
| same_date | 1 | 328.1475 | 3.28E+02 | 28729.52 | 0 | 4.91 |
| water_dif:sed_dif | 1 | 101.5731 | 1.02E+02 | 8892.788 | 0 | 1.52 |
| Residuals | 299145 | 3416.824 | 1.14E-02 | NA | NA | 51.15 |

Full best fit model

| | Df | Sum Sq | Mean Sq | F value | Pr(>F) | Percent Variance Explained |
|---|---|---|---|---|---|---|
| water_dif | 1 | 1475.45 | 1.48E+03 | 136850.79 | 0.0E+00 | 22.09 |
| sed_dif | 1 | 951.56 | 9.52E+02 | 88259.06 | 0.0E+00 | 14.25 |
| geo_dist | 1 | 405.98 | 4.06E+02 | 37655.77 | 0.0E+00 | 6.08 |
| same_date | 1 | 328.15 | 3.28E+02 | 30436.35 | 0.0E+00 | 4.91 |
| water_dif:sed_dif | 1 | 101.57 | 1.02E+02 | 9421.11 | 0.0E+00 | 1.52 |
| water_dif:geo_dist | 1 | 0.83 | 8.26E-01 | 76.60 | 2.1E-18 | 0.01 |
| sed_dif:geo_dist | 1 | 16.57 | 1.66E+01 | 1536.90 | 0.0E+00 | 0.25 |
| water_dif:same_date | 1 | 55.50 | 5.55E+01 | 5147.61 | 0.0E+00 | 0.83 |
| sed_dif:same_date | 1 | 6.67 | 6.67E+00 | 618.97 | 1.7E-136 | 0.10 |
| geo_dist:same_date | 1 | 93.49 | 9.35E+01 | 8671.52 | 0.0E+00 | 1.40 |
| water_dif:sed_dif:geo_dist | 1 | 3.11 | 3.11E+00 | 288.75 | 1.0E-64 | 0.05 |
| water_dif:sed_dif:same_date | 1 | 1.31 | 1.31E+00 | 121.58 | 2.9E-28 | 0.02 |
| water_dif:geo_dist:same_date | 1 | 5.82 | 5.82E+00 | 540.25 | 2.1E-119 | 0.09 |
| sed_dif:geo_dist:same_date | 1 | 2.55 | 2.55E+00 | 236.91 | 1.9E-53 | 0.04 |
| water_dif:sed_dif:geo_dist:same_date | 1 | 5.86 | 5.86E+00 | 543.22 | 4.8E-120 | 0.09 |
| Residuals | 299135 | 3225.105 | 1.08E-02 | NA | NA | 48.28 |

**Table S3**. Sequential analysis of variance of multiple linear regression models. Model 5 represents the model that explains the most variance, while model 6 is the simplified model using only terms that explain >1% of the variance. Water = water depth, sed = sediment depth, distance = geographic distance between sites, environment = binary factor representing N of S Gulf of Mexico, date = factor for sampling date (5 sampling timepoints).

| | Res.Df | RSS | Df | Sum of Sq | F | Pr(>F) |
|---|---|---|---|---|---|---|
| 1. water | 299149 | 5204.1 | | | | |
| 2. water * sed | 299147 | 4150.5 | 2 | 1053.58 | 48860.7 | < 2.2e-16 |
| 3. water * sed * distance | 299143 | 3726 | 4 | 424.51 | 9843.6 | < 2.2e-16 |
| 4. water * sed * environment | 299143 | 3786.5 | 0 | -60.5 | | |
| 5. water * sed * distance * date | 299135 | 3225.1 | 8 | 561.39 | 6508.7 | < 2.2e-16 |
| 6. water + sed + distance + date + water:sed | 299145 | 3416.8 | -10 | -191.72 | 1778.2 | < 2.2e-16 |

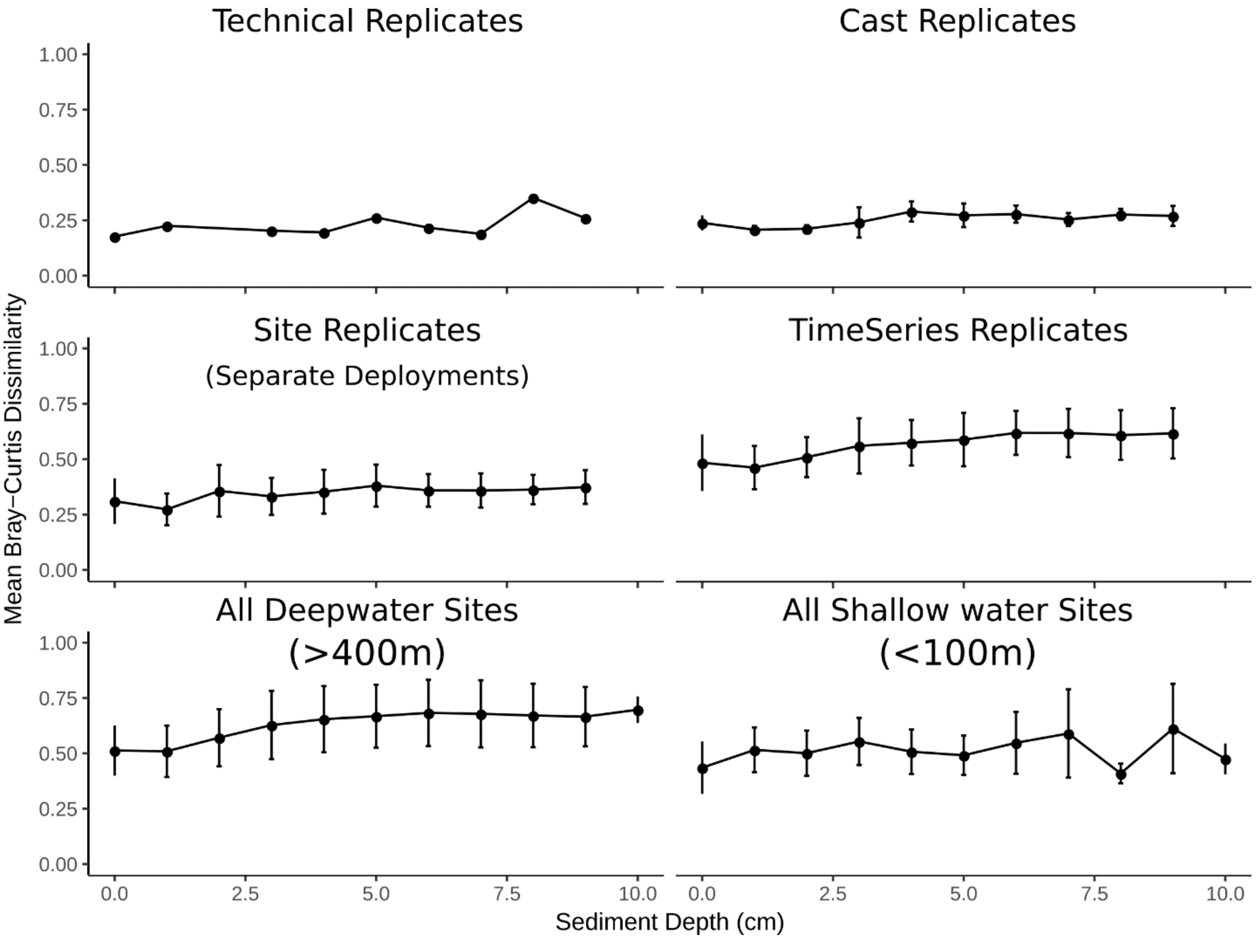


**Figure S1. Changes in microbial community similarity at different levels of replication.** Pairwise Bray-Curtis similarities were calculated for every sample, and samples were binned into 1 cm depth increments to enable cross study comparisons. Average replicate level dissimilarity value is shown in the top right of each plot with the standard deviation. Error bars represent standard deviation about the mean for each sediment depth bin. (A) The same core (AC-5) depth sections were independently extracted and sequenced in duplicate. (B) For three sites, within 1 multicorer cast, triplicate cores were extruded and depth sections were extracted and sequenced. This represents variation within 1m$^2$ of the seafloor. (C) For 10 sites, triplicate multicore deployments were performed, 1 core from each deployment was extruded, and DNA extractions and sequencing was performed on the sections. This represents community variation of tens to hundreds of square meters on the seafloor. (D) Seven sites were sampled over multiple cruises. (E) All sites sampled at water depths > 800m, representing the deep Gulf of Mexico. (F) All sites sampled at water depths less than 100m, representing the shallow Gulf of Mexico.

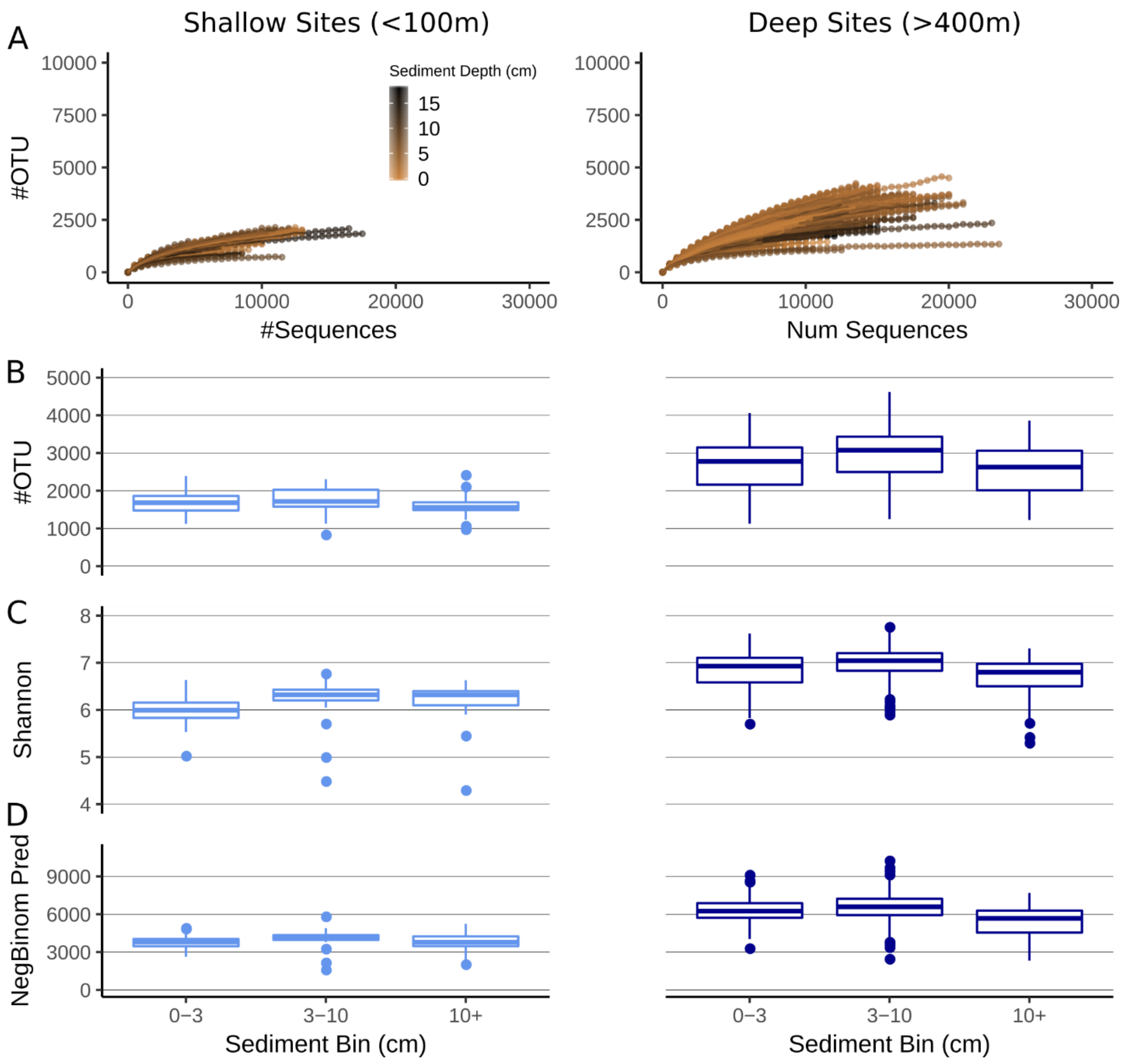


**Figure S2. Microbial alpha diversity metrics from the Gulf of Mexico.** The normalized OTU table that was used for all beta-diversity analyses (Figure 2) was also used here. There are no sites >100 m and <400 m water depth (all samples are included in this figure). (A) Rarefaction curves generated by subsampling at 500 depth intervals, samples are colored based on depth within the sediment (not significant). (B) Observed number of OTUs, (C) Shannon diversity calculated using base e, (D) Estimated number of OTUs using a zero-truncated negative binomial model.

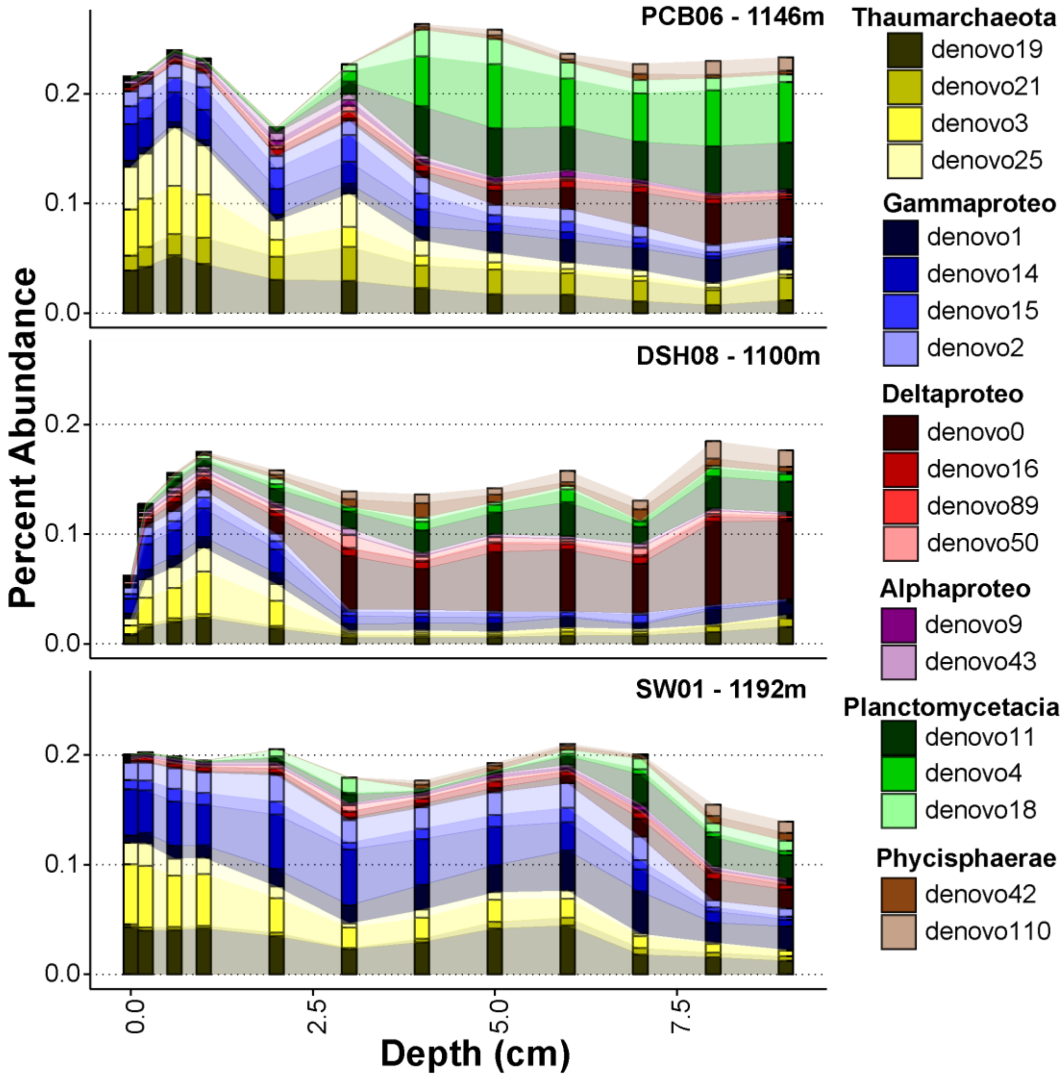


**Figure S3. Core profiles representing an East to West transect sampled at approximately 1100m water depth in the Northern Gulf of Mexico showing population (OTU) level distributions.** Core profiles are arranged from the eastern most site to the western most site.

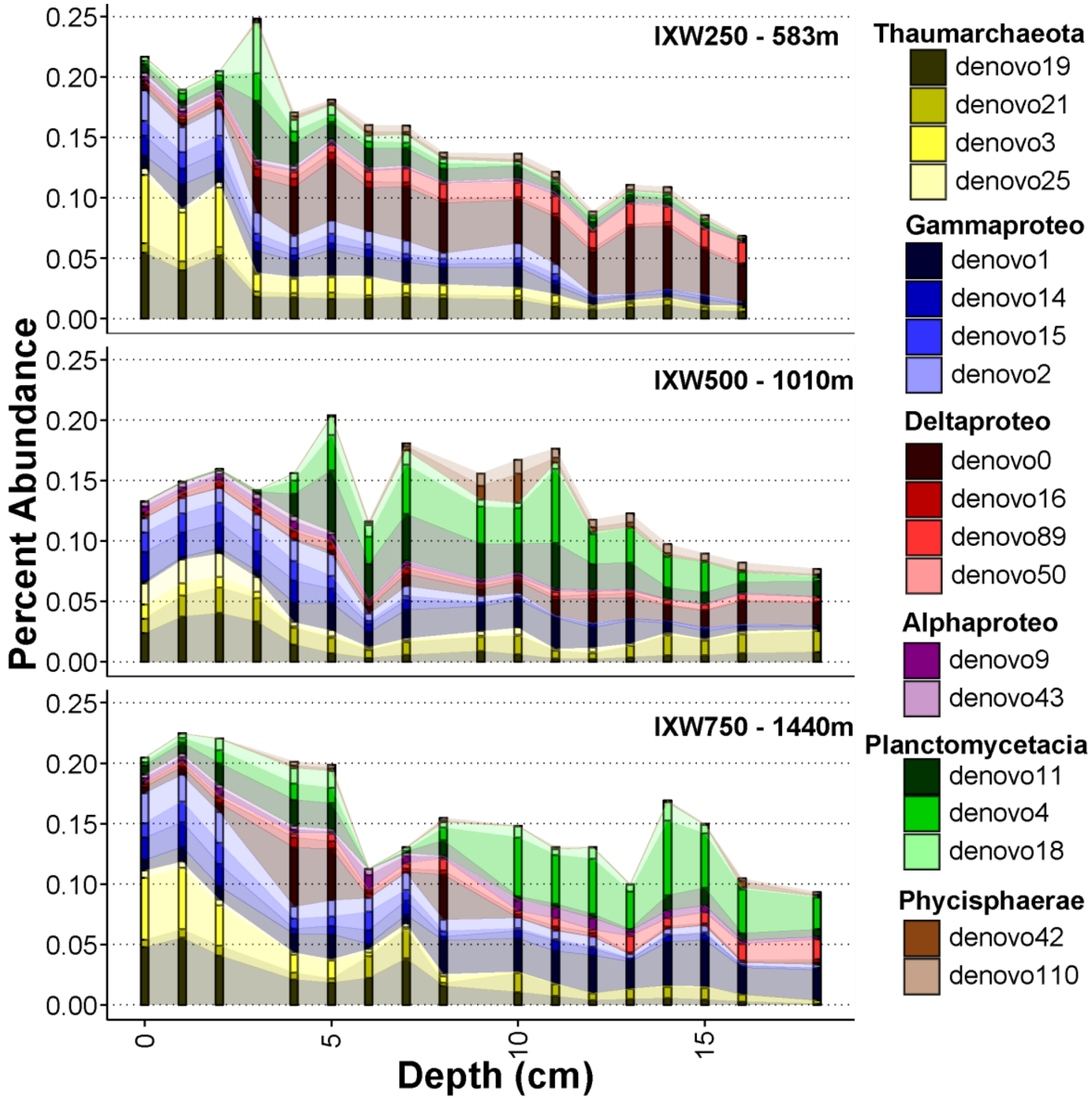


**Figure S4. Core profiles representing a depth transect in the Southern Gulf of Mexico showing population (OTU) level distributions.** Core profiles are arranged with increasing water depth.

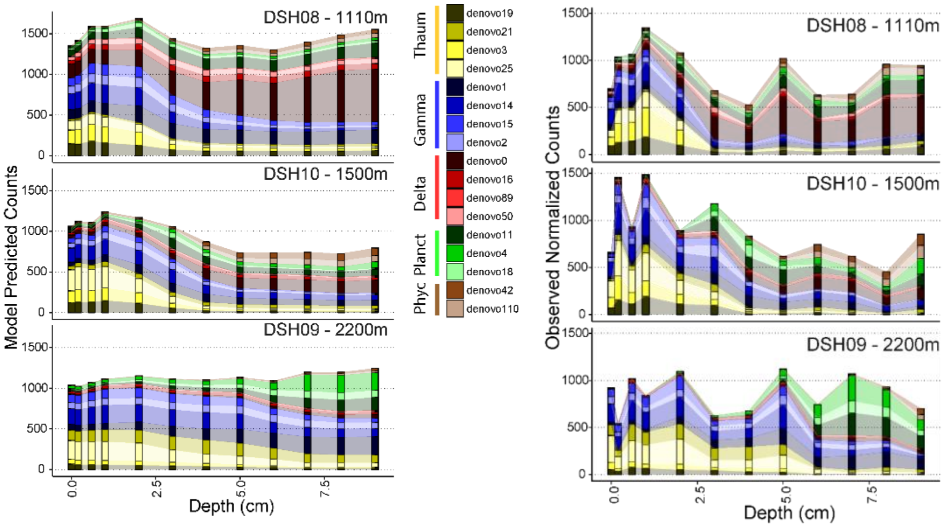


**Figure S5. A comparison of the model predictions for these abundant OTU relative to the observed values shown in Figure 6.**

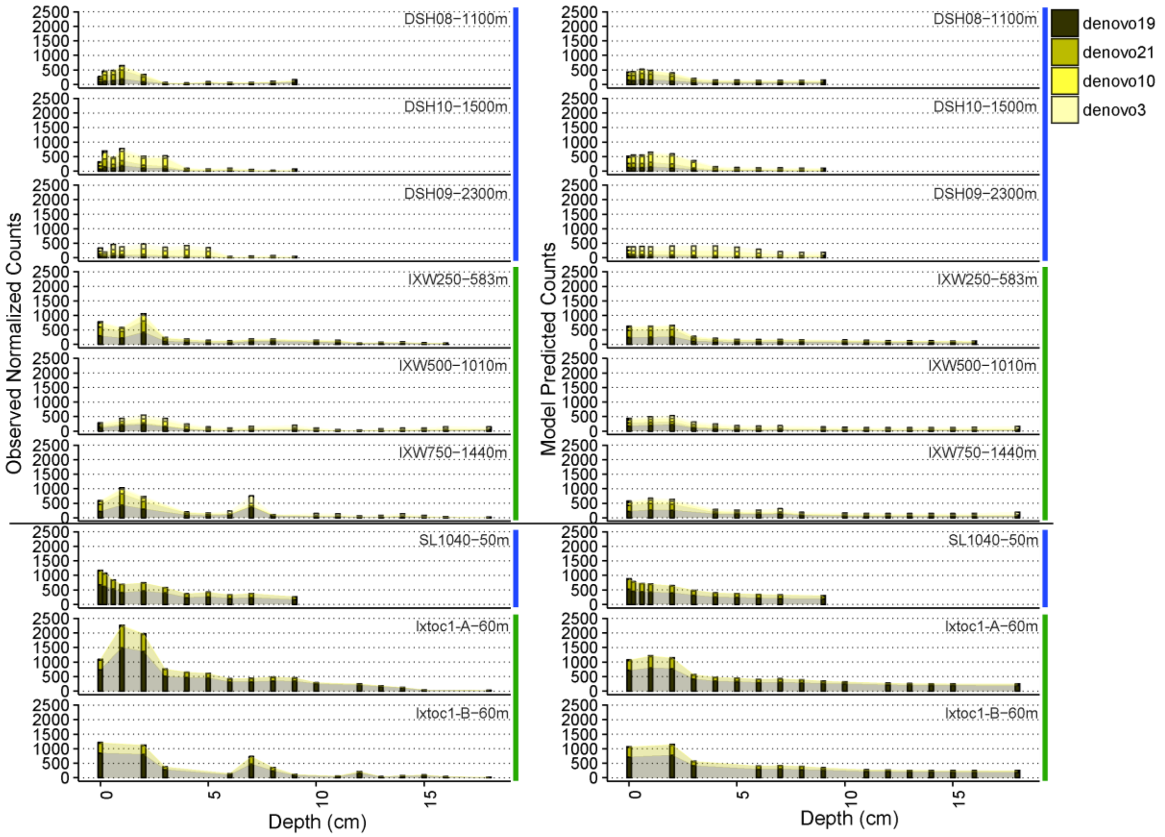


**Figure S6. Biogeography of the dominant Thaumarchaeota OTU.** The left side of the figure shows the observed counts for each of the OTU, while the right side shows the corresponding model predictions. Above the horizontal black line are deep water sites, while those below are shallow water sites. Blue horizontal bars to the right of the figures indicate samples collected in the northern Gulf of Mexico, while green horizontal bars indicates those from the southern Gulf of Mexico.

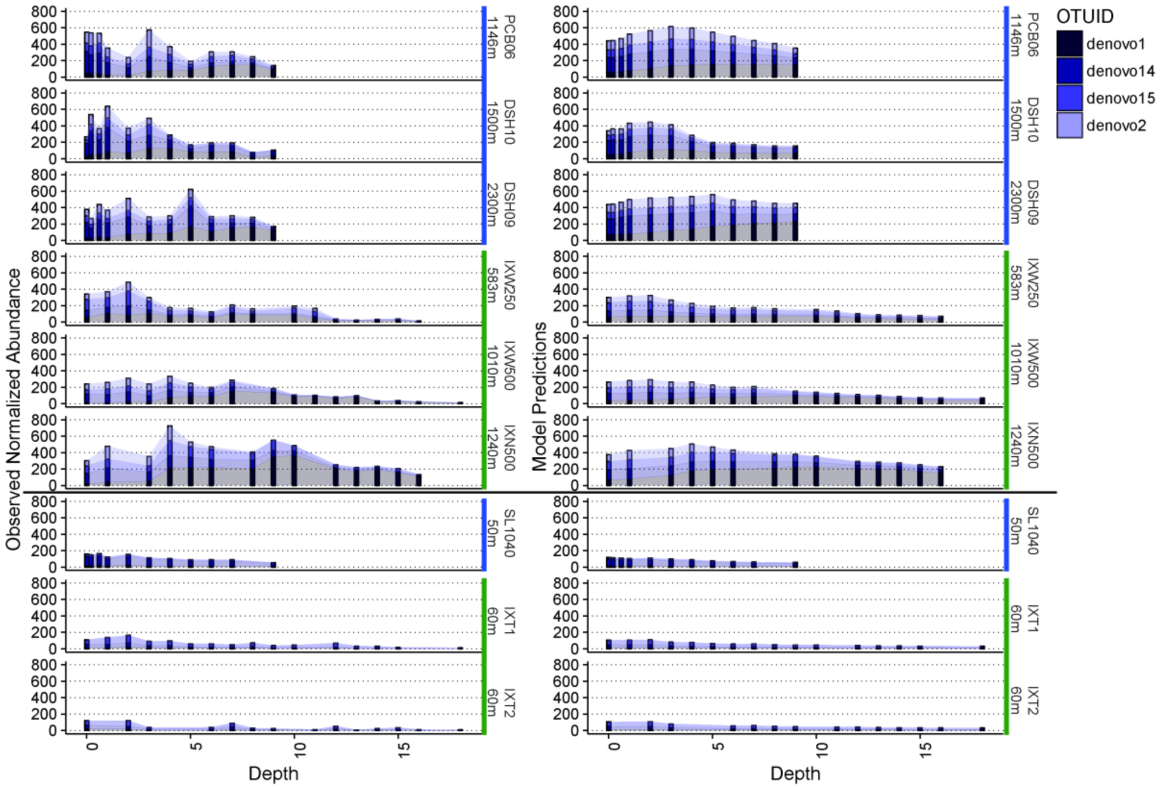


**Figure S7. Biogeography of the dominant Gammaproteobacteria OTU.** The left side of the figure shows the observed counts for each of the OTU, while the right side shows the corresponding model predictions. Above the horizontal black line are deep water sites, while those below are shallow water sites. Blue horizontal bars to the right of the figures indicate samples collected in the northern Gulf of Mexico, while green horizontal bars indicates those from the southern Gulf of Mexico.

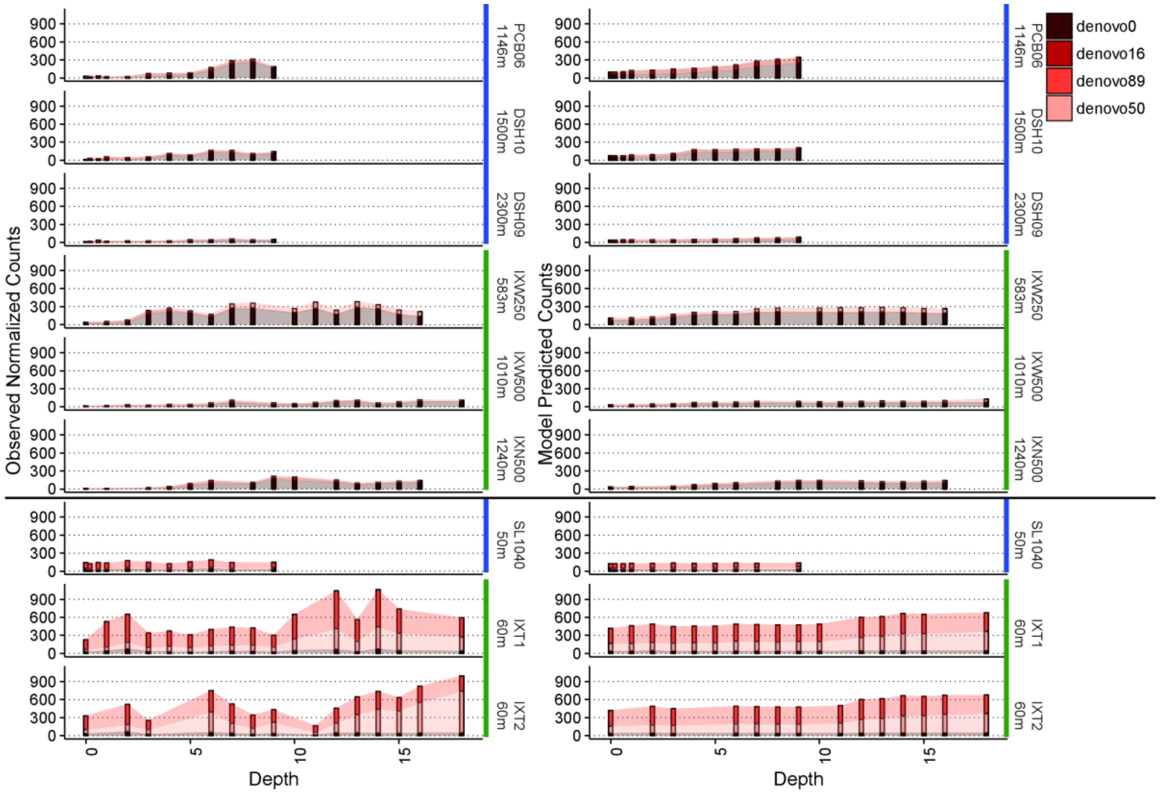


**Figure S8. Biogeography of the dominant Deltaproteobacteria OTU.** The left side of the figure shows the observed counts for each of the OTU, while the right side shows the corresponding model predictions. Above the horizontal black line are deep water sites, while those below are shallow water sites. Blue horizontal bars to the right of the figures indicate samples collected in the northern Gulf of Mexico, while green horizontal bars indicates those from the southern Gulf of Mexico.

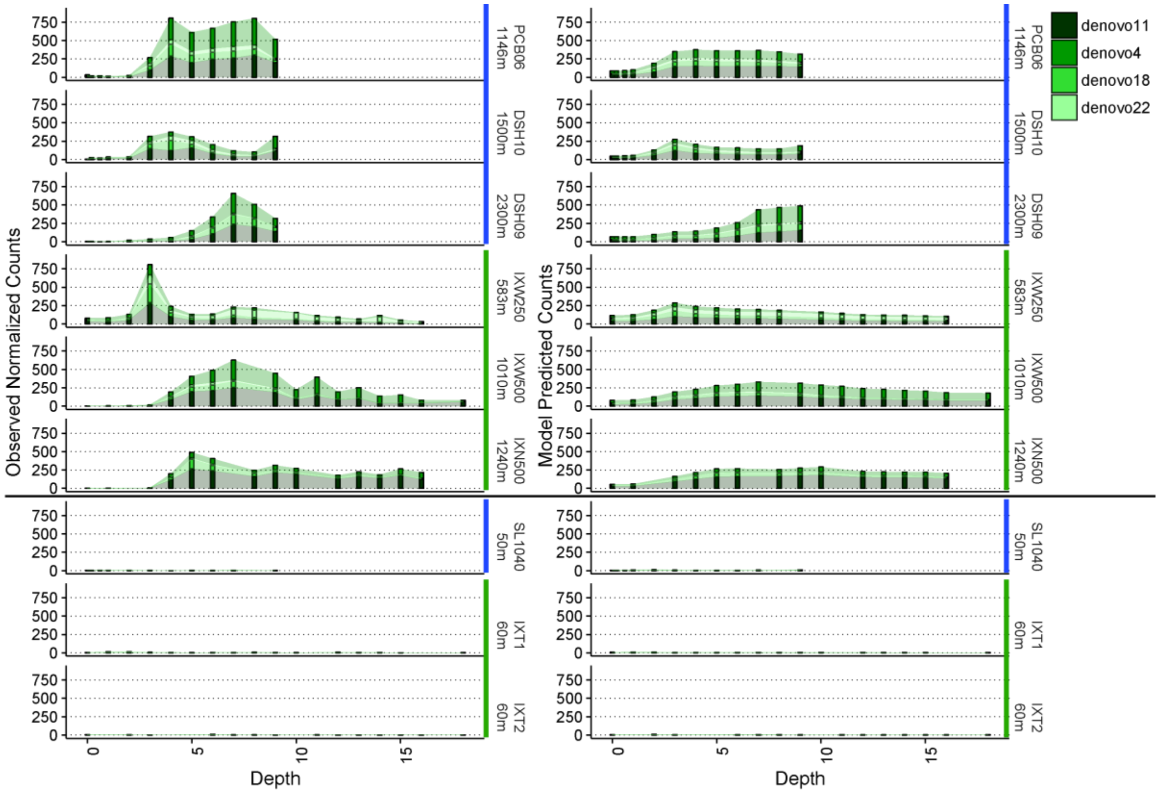


**Figure S9. Biogeography of the dominant Planctomycetes OTU.** The left side of the figure shows the observed counts for each of the OTU, while the right side shows the corresponding model predictions. Above the horizontal black line are deep water sites, while those below are shallow water sites. Blue horizontal bars to the right of the figures indicate samples collected in the northern Gulf of Mexico, while green horizontal bars indicates those from the southern Gulf of Mexico.

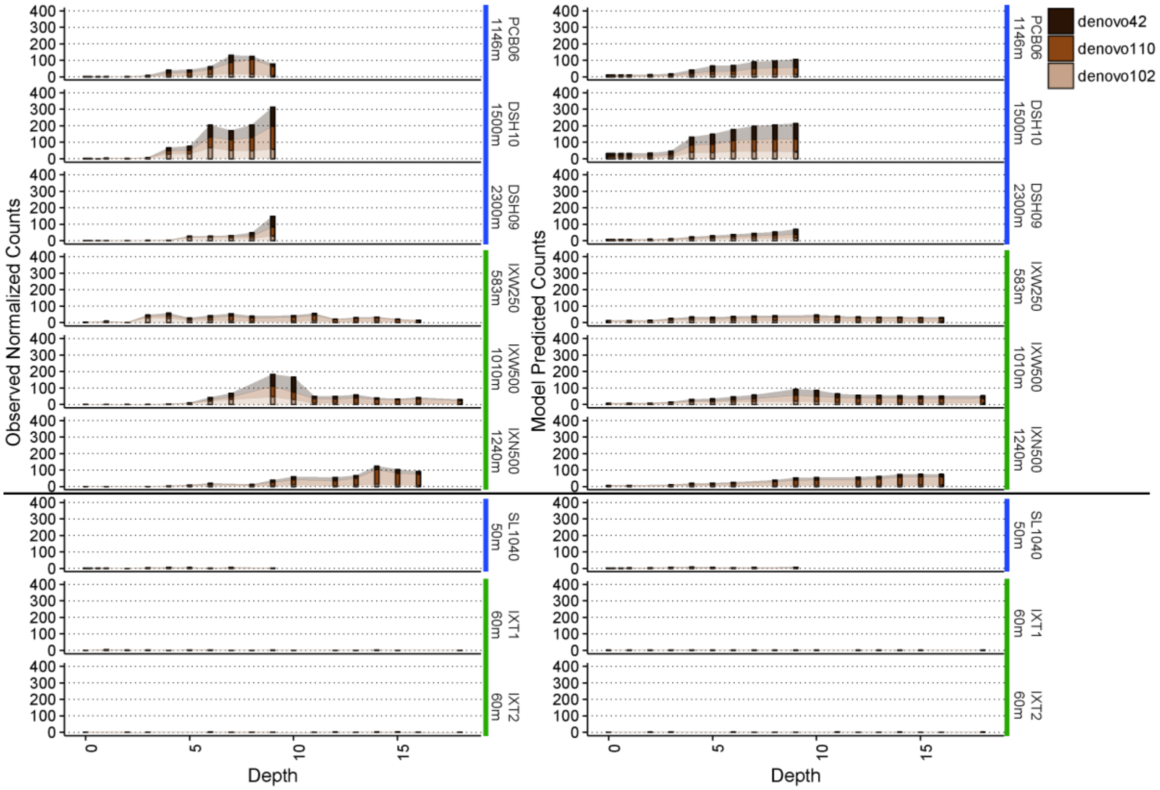


**Figure S10. Biogeography of the dominant Phycisphaerae OTU.** The left side of the figure shows the observed counts for each of the OTU, while the right side shows the corresponding model predictions. Above the horizontal black line are deep water sites, while those below are shallow water sites. Blue horizontal bars to the right of the figures indicate samples collected in the northern Gulf of Mexico, while green horizontal bars indicates those from the southern Gulf of Mexico.

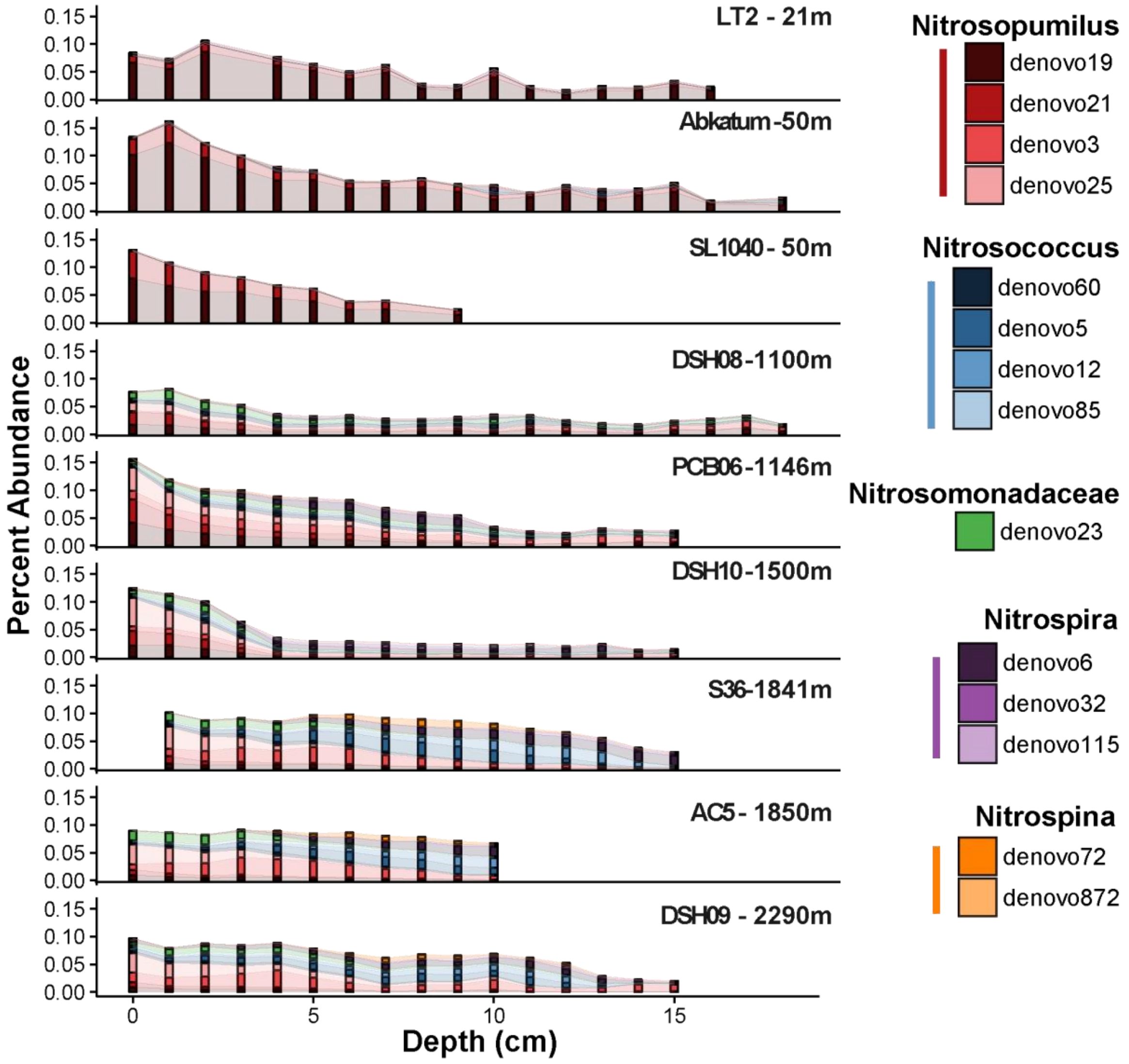


**Figure S11. Biogeographic patterns in putative nitrifying populations throughout the Gulf of Mexico.** All core profiles are organized from the shallowest site at the top to the deepest site at the bottom. Nitrosopumilus, Nitrosococcus, and the Nitrosomonadaceae family are considered as putative ammonium oxidizers, while Nitrospira and Nitrospina are considered here as putative nitrite oxidizers.
